# *ProteoSeeker*: A Feature-Rich Metagenomic Analysis Tool for Accessible and Comprehensive Metagenomic Exploration

**DOI:** 10.1101/2024.10.21.619413

**Authors:** Georgios Filis, Dimitra Bezantakou, Konstantinos Rigkos, Despina Noti, Pavlos Saridis, Dimitra Zarafeta, Georgios Skretas

## Abstract

Metagenomics have served as a key driver of biotechnology advancements through, among others, the identification of novel proteins. The lack of standardized guidelines and benchmarks in the field, however, complicates the selection of appropriate bioinformatics tool ensembles and hinders faster progress. This study introduces *ProteoSeeker*, an automated pipeline designed for accessible and comprehensive metagenomic exploration of whole-genome sequencing data. *ProteoSeeker* identifies proteins within user-defined protein families and uncovers the taxonomy of the host organisms. It is implemented as a command-line tool to facilitate metagenomic dataset analysis for non-expert users, thus enabling facilitated and streamlined protein discovery.

## Background

Approximately 80% of Earth’s microbial diversity remains unculturable, posing a significant limitation in unlocking the functional potential encoded within its genetic content (1). Metagenomic analysis can bypass this bottleneck by providing access to the collective genomic information obtained from environmental samples – referred to as the metagenome (2). The exponential growth of publicly available metagenomic data, driven by advances in DNA sequencing technologies, underscores the critical need for tools to process this vast information reservoir and identify genes encoding biotechnology-relevant proteins, which can significantly drive innovation.

Metagenomic analysis has had a broad impact across various research fields, including microbiome studies, biotechnology, ecology and phylogenetics (3). Additionally, function-driven metagenomics serve as a cornerstone for deducing functional capabilities from genomic/metagenomic data, with the identification of novel genes leading to the discovery of proteins and enzymes with new and/or enhanced functional traits (4). This systematic approach contributes to broadening the spectrum of known valuable biocatalysts and biomolecules, addressing the increasing demand for highly specific and process-compatible biomolecules.

A promising area of metagenomic analysis towards the discovery of new and potentially useful proteins involves studying extremophiles—microbes that thrive in extreme environmental conditions. Adapted to challenging environments, such as high or low temperatures, pH or salinity habitats, extremophiles can harbor unique biocatalysts, known as extremozymes, which possess technologically significant characteristics. These enzymes have demonstrated value in diverse applications spanning agriculture, biochemistry, biomedicine, and beyond (5). Leveraging bioinformatic methods is essential for analyzing the vast available and continuously increasing metagenomic data, thus enabling and accelerating the identification, annotation, and functional prediction of novel extremozymes.

While metagenomic analysis using bioinformatic algorithms is highly advantageous, it is persistently characterized by technological constraints. One of the fundamental processes in metagenomics analysis is generating a “metagenome-assembled genome” (MAG), typically obtained from whole-genome shotgun sequencing. This method involves the fragmentation of the genetic material, enabling the thorough sequencing of all DNA fragments in a sample, even at a single strain level, allowing for detailed functional profiling (6). Analyzing shotgun sequencing metagenomic data presents significant challenges for the researcher, particularly in selecting and combining tools to perform separate activities, such as sequence assembly, binning and annotation. These tasks are complex and time-consuming, requiring a deep understanding of various computational methods and their compatibility.

Well-established platforms, such as MG-RAST, IMG/M and MGnify, are widely recognized choices for performing the main metagenomic analysis steps (7–9). Despite the availability of numerous tools and pipelines that simplify and expedite separate analysis processes, the field lacks standardized guidelines and performance benchmarks, making it challenging to select and curate the most suitable ensemble of bioinformatics tools. Furthermore, many established pipelines incorporate an excess of tools regarding the application of a specific stage or task, making it difficult and time-consuming for a user to decide which is the ideal sequence of tool-utilization to complete the analysis. Thus, there is an unmet need for developing rapid, reproducible, and precise analytical pipelines with the well-defined scope to automate workflows and ensure consistency across studies (10).

To address these challenges, we introduce *ProteoSeeker*, a command-line tool specifically designed for comprehensive functional analysis of metagenomic data from whole-genome sequencing (WGS) and by extension of genomic and proteomic data. The main focus of *ProteoSeeker* is to identify proteins from user-defined protein families or/and uncover the taxonomy of their host organisms. With minimal user input needed, *ProteoSeeker* empowers researchers, including those with limited bioinformatics expertise, to perform protein annotation, protein domain identification, protein family prediction, and taxonomic analyses. As a result, *ProteoSeeker* offers valuable insights into any given metagenomic dataset, streamlining the analytical process and expanding user accessibility in metagenomic research. The introduction of *Proteoseeker* is anticipated to accelerate protein discovery by democratizing further the exploration of metagenomic data. This advancement is poised to make a significant contribution to achieving innovation and sustainability goals across diverse sectors, ranging from agriculture and pharmaceuticals to environmental remediation and beyond, within the ever-evolving landscape of biotechnology.

## Results

### Overview

*ProteoSeeker* is a novel, highly comprehensive pipeline that combines state-of-the-art software for metagenomic analysis of WGS data and it is designed to run effectively on limited computational resources, without hindering scalability. Comprehensibility is achieved through the automation of key processes, such as read preprocessing, contig assembly, gene prediction, putative protein screening and taxonomic analysis. *ProteoSeeker* is designed to run with minimal input. Only an SRA code from the SRA database of NCBI (11,12) or a dataset and in certain cases at least one protein family code have to be provided by the user to initiate the analysis.

The tool offers two main functional modes termed “seek” and “taxonomy”. In both modes, *ProteoSeeker* identifies proteins encoded by predicted protein-coding regions in the contigs assembled from the sequencing reads. In the seek mode, the identified proteins are filtered to select those belonging to the protein family (or families) specified by the user, while in the taxonomy mode, the identified proteins undergo taxonomic classification. Users have the flexibility to select the application of either one or both modes in a single *ProteoSeeker* run.

Each individual tool included in the *ProteoSeeker* pipeline has been specifically selected and optimized to facilitate rapid analysis of the input data. The protein annotation process aims to provide users with insights on which proteins may be suitable for further examination, based on the provided protein annotation. The *ProteoSeeker* pipeline is designed to be highly versatile, enabling users to selectively omit analysis stages based on the scope of their work. Figure 1 illustrates the role of *ProteoSeeker* in the general workflow of protein discovery through metagenomic analysis.

**Figure 1.**
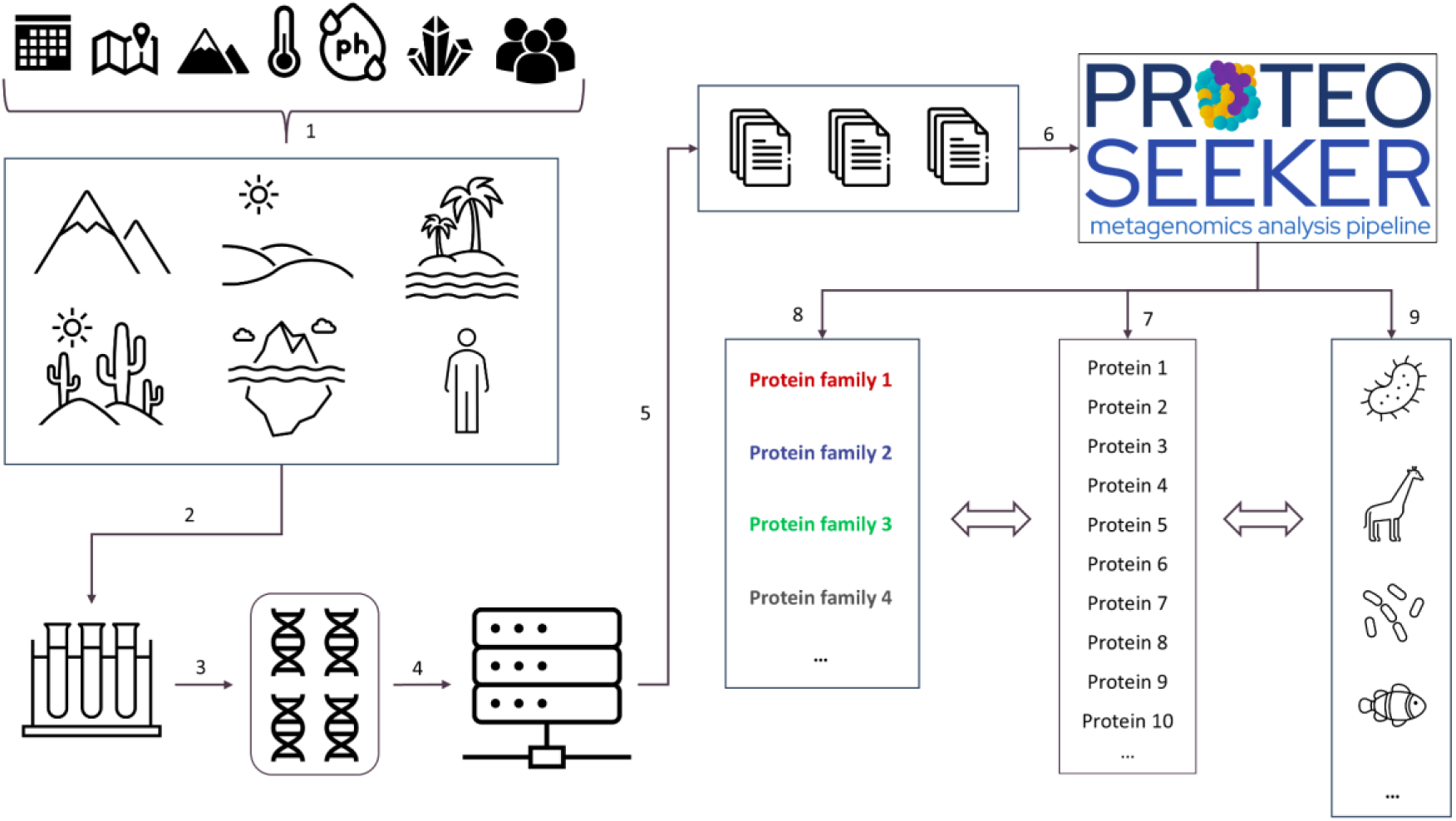
***ProteoSeeker* methodology in the broad context of the standard metagenomics protein discovery pipeline**. **(1)** The targeted sampling environments are selected through defining the specific conditions/origin of interest (organism, temperature, pH, salinity etc.) of the proteins to be discovered and **(2)** the metagenomic sample is collected from the selected environments. **(3)** The DNA of the metagenomic sample is isolated and prepared for sequencing. **(4)** The metagenomic material is sequenced using Next-Generation Sequencing protocols. **(5)** Files containing reads are generated and may be uploaded to open-access databases (as the SRA) alongside their metadata. **(6)** The *ProteoSeeker* user provides an SRA code or a dataset as input. **(7)** *ProteoSeeker* identifies putative proteins from the assembled reads. Users can select to apply the “seek” mode or the “taxonomy” mode or both. **(8)** In the seek mode certain identified proteins are associated with the user-defined protein families based on their corresponding Pfam profiles. This mode is used to uncover novel proteins with targeted functionalities. **(9)** In its taxonomy mode *ProteoSeeker* performs binning and taxonomy classification of the identified proteins.

*ProteoSeeker* offers a multitude of options regarding the user’s desired output. Some of these options control the workflow of the pipeline. As mentioned above, either the seek or the taxonomy mode or both can be applied in a single *ProteoSeeker* run. The seek mode of *ProteoSeeker* includes three types of analysis (“type 1”, “type 2”, “type 3”). Type 1 analysis includes searching for proteins containing at least one domain corresponding to a profile associated with the selected protein families and type 2 analysis includes searching for putative protein hits with E-values lower than a specific score. The hits are acquired by running DIAMOND (13) to screen the putative proteins against a filtered protein database. Type 3 analysis applies both type 1 and type 2 analyses in a single run. The filtering of the protein database has been applied based on the protein names associated with the selected protein families and the provided protein database. Type 1 analysis generates results comprising proteins, which are likely to belong to the selected protein families. Type 2 analysis is more likely to generate results that comprise proteins, which belong to the selected protein families but differ from the characterized family members. These might include proteins that are part of the selected families while sharing distant protein sequences with the other family members or belong to new protein families related to the selected ones. A summary of the results documented in the annotation files generated from a *ProteoSeeker* run after applying both the seek and taxonomy modes, can be found in Additional file 3. The stages of the pipeline in the seek mode of *ProteoSeeker* can be found in Figure 2.

**Figure 2.**
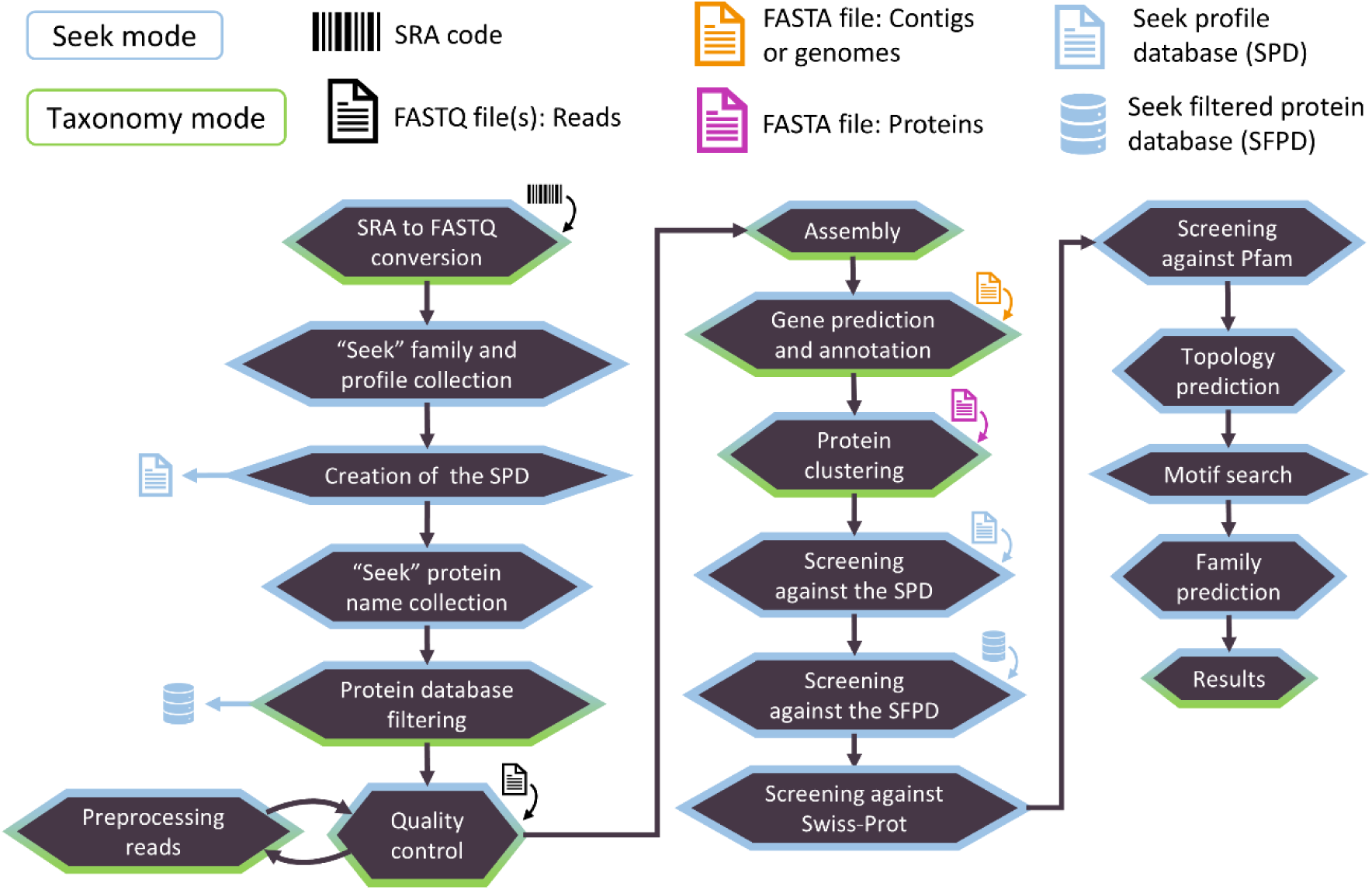
The stages of the “seek” mode of *ProteoSeeker*. *ProteoSeeker* offers the functionalities of the seek mode (blue) and of the “taxonomy” mode (green). Each stage is colored based on the mode it belongs to. The possible types of input for *ProteoSeeker* include an SRA code, reads in FASTQ files and contigs or genomes or proteins in FASTA format. If an SRA code is provided the corresponding SRA and FASTQ files are generated. The “seek” protein families are selected based on the input “seek” family codes and their profiles are collected. The “seek profile database” (SPD) is created. The “seek” protein names of the selected families are collected, and the protein database is filtered based on these names creating the “seek filtered protein database” (SFPD). The reads of the FASTQ files undergo several quality control checks by FastQC. The reads are preprocessed by BBDuk and then are reanalyzed by FastQC. The preprocessed reads are assembled into contigs by Megahit. Protein coding regions (pcdrs) are predicted in the contigs by FragGeneScanRs. CD-HIT is used to reduce the redundancy of the pcdrs. The pcdrs are screened against the SPD with HMMER. Any pcdr with at least one hit based on the latter screening is retained (protein set 1). The rest of the pcdrs are screened against the SFPD with DIAMOND and only those with a hit of low enough E-value are retained (protein set 2). Protein set 1 is screened against the UniprotKB/Swiss-Prot database with DIAMOND. Both protein sets are screened against all the profiles of the Pfam database with HMMER. Topology predictions are performed by Phobius. Motifs provided by the user are screened against each protein. The protein family of each protein is predicted. Annotation files are written.

In addition, the taxonomy mode may be applied through two distinct routes. The “Kraken2 route”, which includes the use of Kraken2 (14,15), Bracken and KrakenTools (16,17) and the “COMEBin/MetaBinner route”, which includes the application of COMEBin (18) or MetaBinner (19). These routes cannot be combined in a single *ProteoSeeker* run, but due to the user’s ability to initiate the run from different pipeline starting points, *ProteoSeeker* may perform taxonomic analysis starting from already assembled contigs. The analysis of Kraken2 and Bracken can be applied based on any Kraken2 and Bracken databases, respectively. The COMEBin/MetaBinner route can be applied with any protein database that includes proteins with a header format that includes information about the organism according to the style used in the headers of the proteins in the non-redundant (nr) database of NCBI (12,20) or the Uniref databases (21–24). Any pre-built or custom-built Kraken2 / Bracken or protein database apart from the default or proposed ones can be downloaded by the user and can be incorporated into the pipeline automatically, requiring as input the paths of these databases in the user’s system. The stages of the pipeline in the taxonomy mode of *ProteoSeeker* can be found in Figure 3.

**Figure 3.**
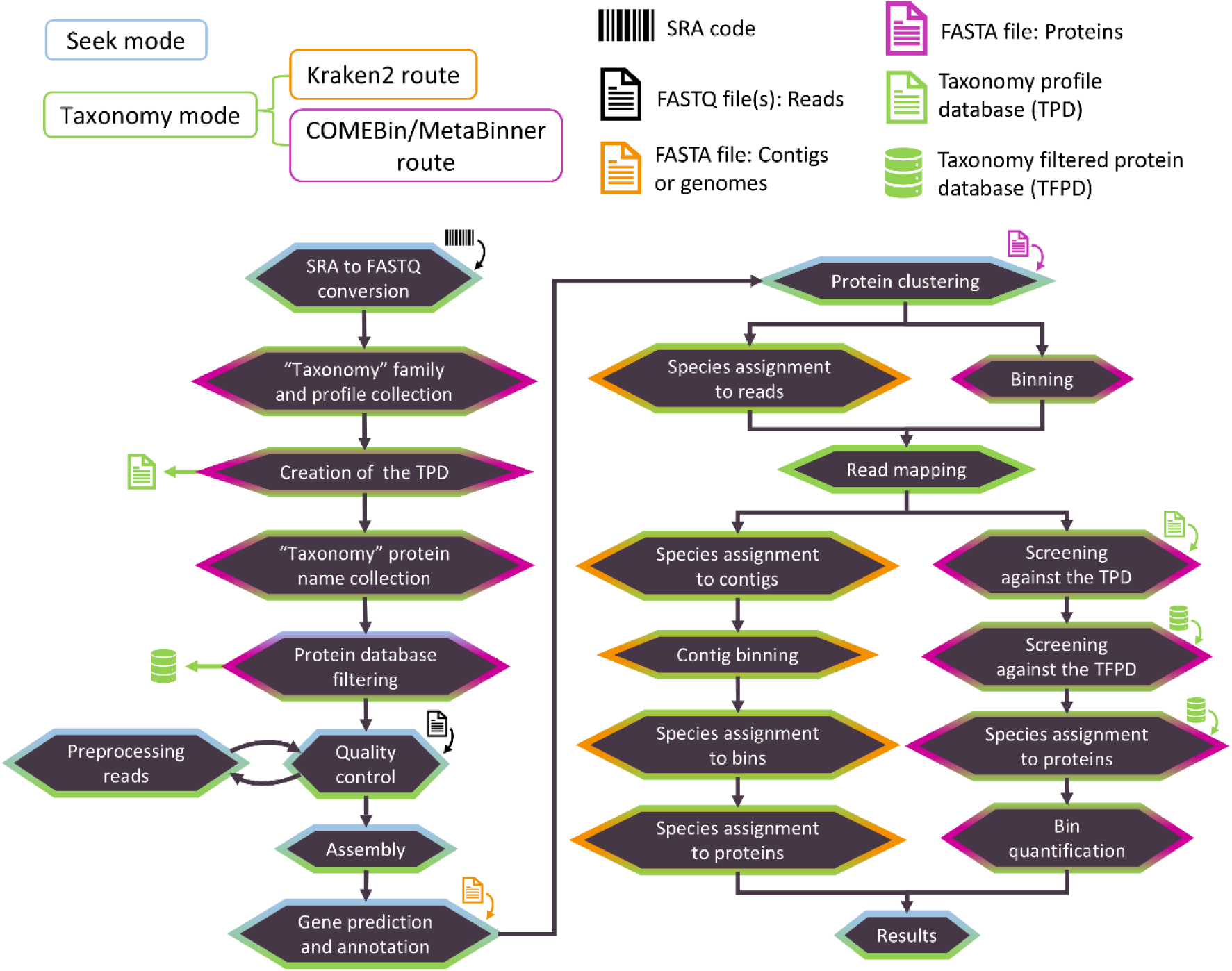
The stages of the “taxonomy” mode of *ProteoSeeker*. *ProteoSeeker* offers the functionalities of the “seek” mode (blue) and of the taxonomy mode (green). The taxonomic analysis is performed by the Kraken2 route (orange) or the COMEBin/MetaBinner route (purple). Each stage is colored based on the mode and route it belongs to. If an SRA code is provided the corresponding SRA and FASTQ files are generated. The “taxonomy” protein families are selected based on the input “taxonomy” family codes and their profiles are collected. The “taxonomy profile database” (TPD) is created. The “taxonomy” protein names of the selected families are collected, and the protein database is filtered based on these names creating the “taxonomy filtered protein database” (TFPD). The reads of the FASTQ files undergo several quality control checks by FastQC. The reads are preprocessed by BBDuk and then are reanalyzed by FastQC. The preprocessed reads are assembled into contigs by Megahit. Protein coding regions (pcdrs) are predicted in the contigs by FragGeneScanRs. CD-HIT is used to reduce the redundancy of the pcdrs. For the Kraken2 route: Species are assigned to the reads based on Kraken2. Bracken then provides the abundances of these species. Bowtie2 maps the reads to the contigs. Species are assigned to contigs. The contigs are binned based on their species. Species are assigned to the bins. Species are assigned to the genes and proteins of the bins. For the COMEBin/MetaBinner route: The contigs are binned based on MetaBinner or COMEBin. Bowtie2 maps the reads to the contigs. The pcdrs are screened against the TPD with HMMER. Any pcdr with at least one hit against the TPD is screened against the TFPD with DIAMOND and any possible hit may provide one or more taxa whose TaxIds and lineages are found through TaxonKit. Taxa are then assigned to bins and to their genes and proteins. Each bin, along with any taxa assigned to it, is quantified. For both routes at the last stage annotation files are generated.

There are multiple options controlling the start and end points of *ProteoSeeker* to facilitate the application of the different seek or taxonomic analysis types and routes, allowing the user to omit prior stages of the pipeline or even subsequent ones. Hence, applying the seek or the taxonomy mode and a route with specific settings in a run can be based on a previous analysis and does not require rerunning the whole analysis. Another time-saving option of *ProteoSeeker* showcasing the versatility of the tool is the option to change the input Kraken2 or Bracken or protein database when reapplying *ProteoSeeker* against a dataset, with the run being initiated after the assembly or gene prediction or the binning process.

*ProteoSeeker* is a Python command-line tool and module, publicly available on GitHub (25) and is also shipped as a Docker image in Docker hub (26). In both forms, an analytical and straightforward installation process manual is provided to the user. All *ProteoSeeker* relevant links, information and instructions can be accessed through the tool’s webpage (27).

### Associating protein families with protein names and Pfam profiles

Specific stages of *ProteoSeeker* depend on the information acquired after processing the UniProtKB/Swiss-Prot (28–30) protein database. This information contains a correspondence of each protein family name with the mean and median length (in number of amino acids) of its members and with the UniProtKB/Swiss-Prot protein IDs, which belong to the members of the protein family, whose lengths were used in computing the median and mean length. Furthermore, the results contain protein family codes and, for each one of these codes, its corresponding protein family name and protein names. Each protein name is accompanied by its frequency rate, which is the number of times this name was found in the names associated with the proteins of the family. Also, the output file contains each protein family name and a corresponding list of Pfam codes for Pfam profiles which correspond to profile hidden Markov models (HMMs) (31,32), accompanied by their frequency and the UniProtKB/Swiss-Prot protein IDs belonging to that family. The list of Pfam codes contains each set of Pfam codes that was computed to have the highest frequency amongst the sets. A set of Pfam codes for a protein consists of each Pfam code corresponding to a domain found in the protein coupled with its frequency. Therefore, it is possible that a protein family may be associated with more than one set of Pfam codes, where all sets have the same frequency. It should be noted that region-specific information about the similarity based on which a protein family is assigned to a protein were corresponded to different protein family names and codes (e.g., similarity based on the notes “In the C-terminal section”, “In the central section” and “In the N-terminal section” for the a protein family name lead to different protein family names and codes) alongside their associated information. The latter approach was followed to avoid cases where a protein might contain a domain or region that is related to a specific protein family, thus the region-specific similarity to that family, while the protein itself might contain additional domains or/and may be part of another protein family. The scripts performing these analyses are available through ProteoSeeker’s publicly available GitHub repository (25). Detailed information about the filtering process of a protein database can be found in Additional file 1: Supplementary Text.

### Experimental validation of the seek mode

We validated the efficiency of the seek mode of *ProteoSeeker* in two distinct discovery expeditions. In both cases, the goal was to discover enzymes with specific functionality and, at the same time, to uncover candidate biocatalysts retaining enzymatic activity at predefined conditions to allow their incorporation in specialized industrial processes. In the first case, we targeted the discovery of thermostable amylolytic α-amylase enzymes with optimal activity at 60-70 °C and neutral pH in order to be incorporated in industrial in-line hydrolysis of starch-rich preparations of infant foods. For this study, we selected as input a number of metagenomic datasets, both in-house and publicly available, with sampling temperatures ranging from 60 to 70 °C and near-neutral pH. Part of these datasets originate from the previously EU-funded project HotZyme (https://cordis.europa.eu/project/id/265933/reporting), in the framework of which numerous biotechnologically valuable novel enzymes have already been discovered (5,33–35). *ProteoSeeker* analysis rapidly led to the identification of proteins containing the targeted protein domains in the analyzed metagenomic datasets. *ProteoSeeker* identified approximately 800,000 protein coding regions from these samples out of which 108 contained at least one domain associated with α-amylases (targeted profile short names: Alpha-amylase, Alpha-amylase_C, alpha-amylase_N, alpha-amyl_C2). To further narrow down the selection and increase the probability of selecting catalytically active protein sequences, the difference in sequence length of each putative protein with the mean length of the members of the “glycosyl hydrolase 13 family” from the UniProtKB/Swiss-Prot database was taken into consideration. In addition to identifying the best match for each protein against the UniprotKB/Swiss-Prot database, *ProteoSeeker* identifies the presence of start and stop codons, as well as the distance of each protein-coding region from the edges of its contig, supporting further the selection of candidate protein sequences for experimental evaluation. This approach highlights the utility of the tool’s output, thereby maximizing the chances of selecting protein sequences likely to exhibit the desired activity.

To showcase the efficiency of *ProteoSeeker*, we evaluated the annotation information provided by the tool and three proteins exhibiting low similarity to proteins of the UniProtKB/Swiss-Prot protein database were selected. Their DNA sequences were codon-optimized for recombinant production in *Escherichia coli*, overexpressed and purified. Their biochemical characterization revealed that all selected proteins (AL_6, AL_15, AL_17) exhibit amylolytic activity within the targeted temperature and pH range (Additional file 1: Figure S1).

Subsequently, *Proteoseeker* was applied for the targeted discovery of carbonic anhydrases (CAs) for industrial CO_2_ capture applications. The discovery of such biocatalysts is rather challenging as the enzyme is intended for industrial CO_2_-capture applications and required to perform under hot potassium carbonate (HPC) conditions, which include temperatures above 80 °C and pH of 11.5. These stringent biochemical requirements necessitated exquisite thermostability and performance under high alkalinity, which are exceptionally rare characteristics for natural proteins. By utilizing these criteria to guide metagenomic dataset selection, the utilization of *Proteoseeker* led to the discovery of CA-KR1 (36), a novel CA exhibiting unprecedented stability and tailored biochemical characteristics that meet the demands of industrial CO_2_ capture pipelines. The discovery of CA-KR1, which, to the best of our knowledge, is the most stable CA known to function under HPC conditions, highlights the significant potential of *ProteoSeeker*. To identify CA-KR1, *ProteoSeeker* analyzed 28 metagenomic datasets, which were selected based on their annotated sample collection temperatures and pH. The threshold of minimum temperature for the selection of the datasets was 80 °C to accommodate industrial application conditions. Approximately 100,000 protein coding regions were predicted during the runs of *ProteoSeeker* for the 3 datasets corresponding to the highest collection temperatures based on their environmental samples. From these protein coding regions *ProteoSeeker* predicted 31 proteins with domains corresponding to profiles associated with CAs through its type 1 analysis in the seek mode. Through the type 2 analysis of the seek mode, *ProteoSeeker* detected approximately 76 proteins having at least one hit with an E-value equal to or below 1e-70 through DIAMOND against the “seek filtered protein database” (SFPD). Manual analysis of the output information provided by *ProteoSeeker,* including evaluation of the output data as mentioned above (mean protein length, start/stop codon presence, distance of the protein coding region from the edges of its contig, etc) led to the selection of 9 proteins to be experimentally tested. The coding DNA sequences of the latter proteins were cloned and expressed in *E. coli*. Out of them, three proteins, CA-KR1 (36), CA_89 and CA_201 (unpublished) were produced successfully and exhibited CA activity, showcasing the significant biotechnological potential of *ProteoSeeker*. The gene sequence of CA-KR1 has been submitted to GenBank (37–39) under the accession number: “BK065798”. Further information regarding the biochemical characterization of CA-KR1 can be found in the work of Rigkos *et al.* (36).

### Evaluation of the taxonomy mode

To analyze and evaluate the different taxonomy routes of *ProteoSeeker*, 19 SRA files corresponding to “gold standard” samples of artificial (simulated) nature and samples originating from cultures of known species compositions, were analyzed as described in the work of Poussin *et al.* (40). Each gold standard dataset contains a specific number of species. Each of these abundances approximates one of the following numbers: 10, 40, 120, 500, 1000. Hence, each sample has been categorized based on the closest of these numbers to its abundance, forming 4 groups of 3 samples each, and 1 group of 7 samples. The latter group corresponds to the group of 10 species, and it includes 4 samples originating from ZyMoBIOMICS (40). Each group contains three samples of artificial origin, one with no bias, one being AT-rich biased and one being GC-rich biased in terms of DNA base content (Additional file 1: Table S1).

Initially, each of the samples above was downloaded and processed from *ProteoSeeker* using its SRA code as input. Then, *ProteoSeeker* analyzed each sample through five different runs, each time utilizing a different taxonomy route or protein database or Kraken2 / Bracken database. As a result, three runs used the Kraken2 route based on the Kraken2 / Bracken Refseq indexes of the Standard-8, Standard-16 and Standard collections (41), which are databases of sizes of 8 GB, 16 GB and 77 GB, respectively. For each of these runs, Bracken was applied to estimate the abundances of the species predicted by Kraken2 by filtering out each species with reads less than 10 and by targeting the species level. A “taxonomy filtered protein database” (TFPD) was created from the nr database in a *ProteoSeeker* run, independently of the evaluation analysis, and was based on protein families associated with RNA polymerases (“RNApol TFPD”). One run used the COMEBin/MetaBinner route, utilizing COMEBin and was based on the RNApol TFPD. One run used the COMEBin/MetaBinner route, utilizing MetaBinner and was based also on the RNApol TFPD. Each run generated a set of predicted species accompanied by their abundances and relative abundances, corresponding to the sample being analyzed in the run. The Kraken2 route allows for the use of multiple threshold values for the filtering step of the species after Bracken has been applied. For the purposes of the evaluation, the threshold values applied were 0.01%, 0.1%, 1.0%, 5.0%, 100, 500, 1000 and the threshold automatically computed by *ProteoSeeker* based on the Shannon index (40,42,43) value for the case of “non-gut” and “gut” samples according to the methodology described by Poussin *et al.* (40). The Shannon index was computed based on the KrakenTools (17). In total, nine filtering thresholds were used for the evaluation. The annotation files contain binning and taxonomy information based on the species remaining after filtering the species of Bracken based on a specific filtering threshold (can be user-defined). Bracken was applied with read length equal to 100, which is the closest value to the average length of the reads in each sample, and with threshold equal to 10 reads as proposed by the protocol described by Lu *et al*. (17). The COMEBin/MetaBinner route allows for more than one species to be associated with a bin, in which case they also share the same abundance and relative abundance. The Shannon index and the value of the filtering threshold computed automatically for non-gut and gut samples for each Kraken2 database and each sample can be found in Additional file 4. The metrics selected to evaluate the results generated by *ProteoSeeker* in each of the 5 analysis cases compared to the gold standard results provided by each of the 19 gold standard samples were the “True Positive (TP)” hits, “False Positive (FP)” hits, “False Negative (FN)” hits, “Sensitivity”, “Precision”, “Accuracy”, “F1 Score”, “Jaccard Index” and “L1 Norm” (40,44–46).

The evaluation of the taxonomy mode took advantage of the stage initiation options of *ProteoSeeker*. Before any of the runs were executed, individual procedures of *ProteoSeeker* were performed individually - the creation of the profile databases, the filtering of the protein database and the collection and processing of each SRA dataset. These operations were performed separately to showcase that the procedures related to processing an SRA code and creating the profile and filtered protein databases can be performed once and then can be used repeatedly by any subsequent *ProteoSeeker* run for either the seek or the taxonomy mode. Hence, the SRA dataset is downloaded and processed once and then each subsequent *ProteoSeeker* run recognizes the presence of the SRA sample locally and utilizes it directly (similarly for the profile and filtered protein databases). The creation of the profile and filtered protein databases by *ProteoSeeker* was based on nr by utilizing 16 central processing units (CPUs) and took approximately 38 min to complete. Processing an SRA dataset involves converting the SRA file to one or more FASTQ files. *ProteoSeeker* examines whether the output FASTQ files are paired-end or single-end or both and proceeds accordingly.

The first run of each sample analysis used the taxonomy route of Kraken2 and the Standard-8 collection as a database. The following 2 runs of the same route initiated *ProteoSeeker* after the stage of gene prediction and used the Standard-16 and Standard collections as databases respectively. Then, the COMEBin/MetaBinner route was used, with COMEBin, where *ProteoSeeker* started the analysis after the stage of gene prediction. The latter procedure was followed also for the COMEBin/MetaBinner route, with MetaBinner. The taxonomy routes, route-specific tools and databases used in the evaluation can be found in Additional file 1: Table S2. All runs of *ProteoSeeker* were performed in an Ubuntu 24.04 LTS system with 124 GBs of RAM and 32 CPUs available. *ProteoSeeker* was run with 24 CPUs, meaning that every tool or custom process run in the pipeline of *ProteoSeeker* utilized at most 24 processes or threads when such an option was available for that tool or custom process. In the Kraken2 taxonomy route, Kraken2 was not applied with memory mapping and in the COMEBin/MetaBinner taxonomy route, COMEBin utilized an NVIDIA GeForce RTX 4090 graphics processing unit (GPU). In addition, only contigs with a length higher than 1000 nucleotides were used in the binning processes of COMEBin and MetaBinner. The parameter files and test scripts used to perform the evaluation can be found in the GitHub repository of *ProteoSeeker* which is publicly available (25).

The results were initially processed according to all Bracken filtering thresholds applied for the Kraken2 taxonomy route and all Kraken2 databases used in the runs (Additional file 2). The best scoring combinations of Kraken2 databases and filtering thresholds based on each metric, were used in evaluating the results collected from the COMEBin/MetaBinner taxonomy route with COMEBin and MetaBinner based on the nr protein database (Figure 4 and Additional file 1: Figures S2 and S3). These combinations include the database of the Standard-8 collection with the filtering threshold of 5.0% and the filtering threshold computed for non-gut samples, the database of the Standard-16 collection with the filtering threshold of 5.0% and the database of the Standard collection without a filtering threshold and with the filtering thresholds of 100 and of 5.0%. The best score for a metric is equal with the highest frequency combination(s) based on their scores across the 19 gold standard samples for that metric. The best score is either the highest (for sensitivity, precision, accuracy, F1 score, Jaccard index, true positive hits) or the lowest (for L1 norm, false positive hits, false negative hits) score depending on the metric. A combination having the best score in a number of the 19 samples would get a frequency equal to that number. The frequency of best scores of each combination for each metric across all 19 samples is found in Additional file 2.

**Figure 4.**
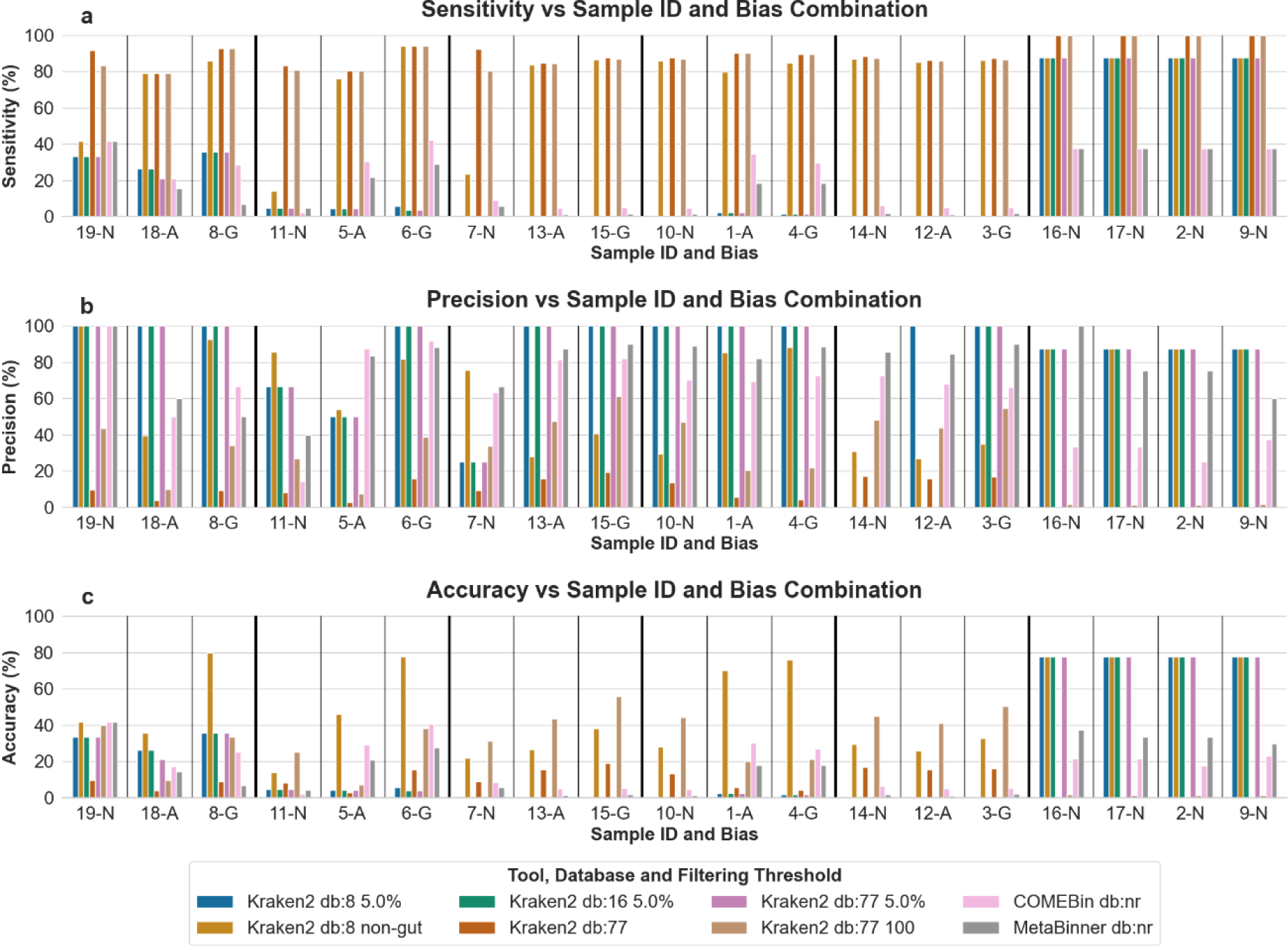
Sensitivity (a), precision (b) and accuracy (c) of the taxonomy mode evaluation results, with the samples sorted based on their species-abundances and biases. The results were acquired by the *ProteoSeeker* runs for the selected combinations of Kraken2 databases and filtering thresholds plus the COMEBin/MetaBinner taxonomy methods for each sample of the 19 gold standard datasets. The selected combinations regarding the Kraken2 taxonomy route include the database of the Standard-8 collection with the filtering threshold of 5.0% (“Kraken2 db:8 5.0%”) and the non-gut filtering (“Kraken2 db:8 non-gut”), the database of the Standard-16 collection with the filtering threshold of 5.0% (“Kraken2 db:16 5.0%”) and the database of the Standard collection without a filtering threshold (“Kraken2 db:77”) and with the filtering thresholds of 5.0% (“Kraken2 db:77 5.0%”) and of 100 (“Kraken2 db:77 100”). The COMEBin/MetaBinner route was applied through COMEBin with the nr protein database as the filtering target (“COMEBin db:nr”) and through MetaBinner with the nr protein database as the filtering target (“MetaBinner db:nr”). The samples are sorted into groups of species-abundances. Samples 19, 18, 8 for 10 species from simulated reads, samples 11, 5, 6 for 40 species, samples 7, 13, 15 for 120 species, samples 10, 1, 4 for 500 species, samples 14, 12, 3 for 1000 species and samples 16, 17, 2, 9 for 10 species from cultures. The letters “N”, “A” and “G” on the labels stand for “No bias”, “AT-rich bias” and “GC-rich bias” respectively.

The time needed to apply Bracken and filter the report of Bracken is negligible compared to the time needed for the analysis to take place (< 1 min) as shown also by the work of Lu *et al.* (17). The execution time of *ProteoSeeker* for the taxonomy evaluation did not include the application of Bracken and the filtering process based on its results, though it included applying the filtering thresholds directly to the results of Kraken2. The time difference of filtering the results of Bracken instead of Kraken2 is negligible. The execution time was also based on the post-classification taxonomy-related processes (e.g., binning, annotation file generation) of the species filtered from the output of Kraken2 based on a specific filtering threshold. Therefore, while the reported execution times do not include the Bracken analysis and are based on selecting the species from the Kraken2 output, they form a solid basis to compare the time efficiency of the different combinations of taxonomy routes and databases and of their individual stages in the pipeline. In general, Bracken is applied rapidly, the filtering thresholds are applied to its output in an identical way as to the output of Kraken2 and the filtered species consist of an unknown set of species *a priori*, which is determined based on the abundance of species and one specific filtering threshold. Consequently, the computed execution times are a representative example of how the execution time of *ProteoSeeker* behaves in each stage and how it differs between the two taxonomy routes and between the different databases used in the Kraken2 route based on the taxonomy evaluation (Figure 5).

**Figure 5.**
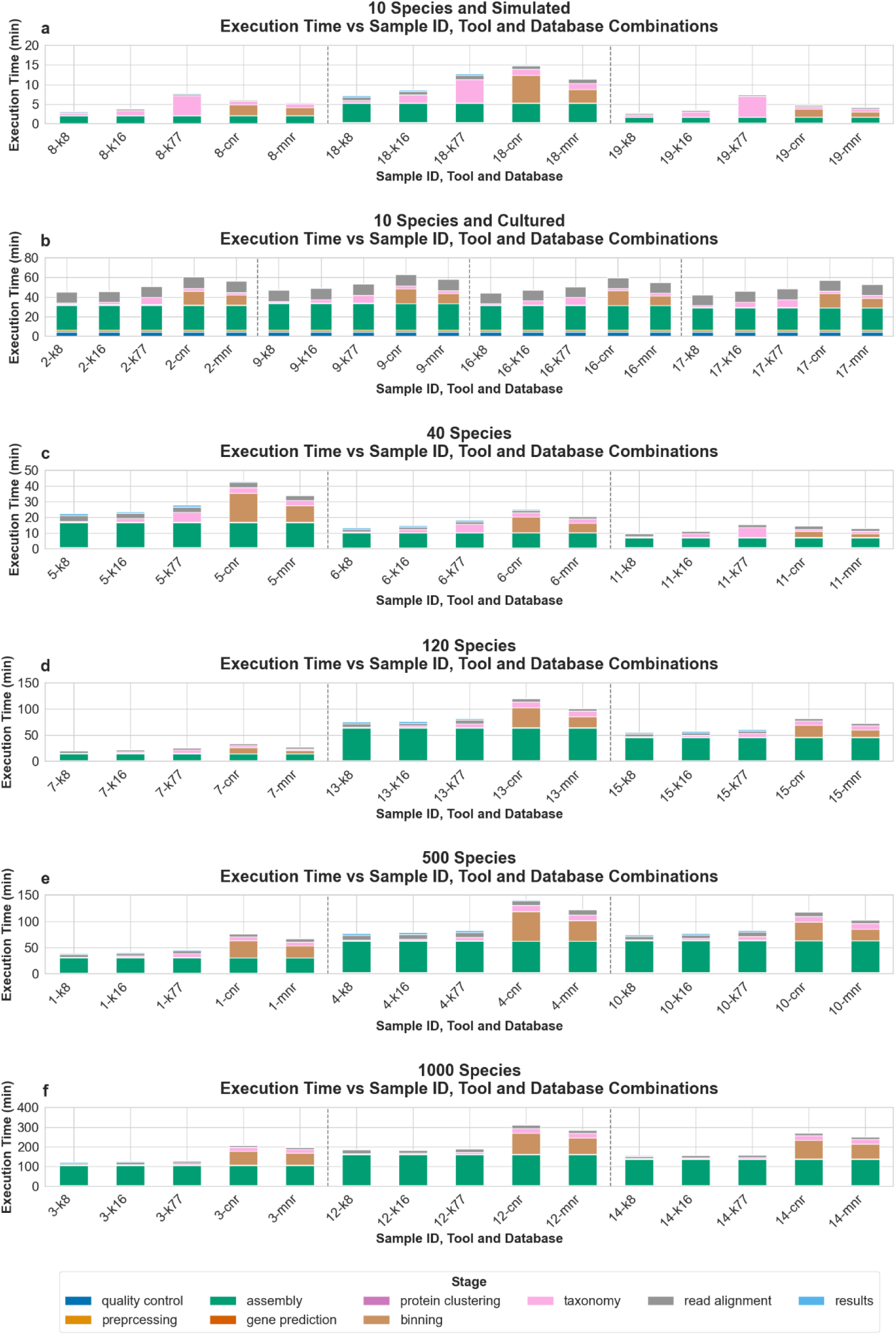
Stage-specific execution times of *ProteoSeeker* runs for each sample, tool and database. Each bar is labeled as “sample ID-tool-database”. For each sample the times related to the runs with the Kraken2 taxonomy route and databases of the Standard-8 collection (“k8”), Standard-16 collection (“k16”) and Standard collection (“k77”), plus the COMEBin/MetaBinner taxonomy route with COMEBin (“cnr”) and MetaBinner (“mnr”) with the nr database are shown. The stage of creating the profile and filtered protein databases and the stage of collecting and processing each SRA sample are excluded from the time analysis shown in the plots. The stages up to and including the stage of gene prediction share the same execution times for all methods of each sample. The plots have been divided into 5 categories, one for the group of 10 species which originate from simulated reads (**a**), one for the group of 10 species originating from cultures (**b**), one for the group of 40 species (**c**), one for the group of 120 species (**d**), one for the group of 500 species (**e**), and one for the group of 1000 species (**f**).

The results of the COMEBin/MetaBinner taxonomy route for each of the two binning tools may include multiple species associated with one bin ID and one relative abundance. Based on the statistical analysis of the results, which is not included in the pipeline of *ProteoSeeker*, the species associated with bin IDs, which numbered more than 1, were discarded. This allowed us to make a more stringent selection of the species that were identified and associated with bin IDs and thus each sample. A species solely associated with more than one bin ID, receives a relative abundance equal to the sum of the relative abundances of its associated bin IDs. This type of filtering for the species predicted by the COMEBin/MetaBinner route can be performed through an additional script available in the GitHub repository of *ProteoSeeker* (25). The same script, apart from generating a new filtered species report can also generate a new TXT annotation file updated based on the filtered species and merged bin IDs. The bin IDs of the same species each of which are assigned only one species are “merged” as their relative abundances for their common species are summed up.

In order to acquire comparable total execution times for the different *ProteoSeeker* runs during the evaluation of the taxonomy mode, specific additions to the execution times of specific runs had to be made due to the differential stage initiation process of *ProteoSeeker* between the runs. Therefore, when computing the stage-specific and total execution time of the *ProteoSeeker* runs, the execution time for the initial stages of the pipeline was acquired by the first run of the tool which was the run for the taxonomy route of Kraken2 with the Standard-8 collection as a database. These initial stages refer to the stages of read quality assessment, read preprocessing, read assembly and gene prediction. Therefore, the latter stages have the same execution times between all runs. The total and stage-specific execution times of *ProteoSeeker* gave us an insight on which stages act as the bottleneck in the analysis and formed the basis of understanding on how *ProteoSeeker* may be improved in the future in terms of speed (Figure 5).

In addition, the execution time of *ProteoSeeker* was analyzed based on the size of the dataset under analysis and the number of species in that dataset for each combination of taxonomy method and database (Additional file 1: Figures S4 and S5). Specifically, for the case of the species abundance, the mean time was computed for each sample category of species abundances (10, 40, 120, 500 and 1000) as well as for the 2 subgroups of the samples with 10 species which originate from simulated reads and cultures respectively.

### Taxonomy mode evaluation for proteins of expected protein families

The taxonomy mode of both routes was also applied to the FASTQ samples, in which the novel ALs and CAs enzymes described before were identified. Based on our search for novel ALs and CAs with specific biochemical characteristics, five enzymes were experimentally profiled. Two of these enzymes (AL_6, AL_15) were identified by running *ProteoSeeker* on a file with contigs, where the taxonomy routes cannot be applied. Therefore, each of the other three enzymes (CA-KR1, CA_201, AL_17) was used to compare the taxonomy results between the Kraken2, the COMEBin/MetaBinner taxonomy routes of *ProteoSeeker* and the taxonomy classification of its best hit based on the results acquired by running BLASTP from the online available blastp suite of NCBI to screen the enzyme against the nr database based on the default parameter values (47). The results acquired from the *ProteoSeeker* runs and the blastp suite can be found in Additional file 1: Table S3. In both cases of mining for ALs or CAs, the protein families selected for the taxonomy route of COMEBin/MetaBinner were associated specifically with the targeted protein family. *ProteoSeeker* was run on the same system as the one used to run the taxonomy evaluations of the gold standard samples. While using the seek mode of *ProteoSeeker* to search for proteins in the family of β-CAs, the taxonomy route of COMEBin/MetaBinner was applied based on different protein families (e.g., α-, β- and γ-CA families) and protein names associated in general with CAs. Similarly, while searching for proteins in the protein family of α-amylases, the taxonomy route of COMEBin/MetaBinner was applied by searching for different protein families and protein names associated in general with ALs. The taxonomy route of COMEBin/MetaBinner was applied using COMEBin and MetaBinner with the nr database as the protein database to be filtered and the taxonomy route of Kraken2 was applied based on the databases of the Kraken 2 / Bracken Refseq indexes of the Standard-8, Standard-16 and Standard collections. For either one of the routes no filtering threshold was applied to the predicted taxa.

## Discussion

*ProteoSeeker* identifies proteins with targeted functionalities by automatically processing metagenomic data from WGS or genomic or proteomic data, while offering taxonomy prediction of the organism(s) associated with each identified putative protein*. ProteoSeeker* has been implemented as a command-line tool and module developed in the Python programming language, is publicly available in GitHub (25) and it is also shipped as a Docker image available publicly in Docker hub (26). The tool’s functionalities are divided into two main modes of function, each mode including its own types and routes of analysis. The functions of the tools in the pipeline and the analysis course can be controlled by the user through the options offered by *ProteoSeeker*. The user can run *ProteoSeeker* with minimal requirements for input, by providing only an SRA code or an input FASTQ/FASTA file and when required selecting the protein family (families) of interest. The output of *ProteoSeeker* is also user-input dependent allowing the results to be documented in respect to whether the user prefers to include the information collected by the seek or the taxonomy mode or both.

In the taxonomy route of COMEBin/MetaBinner, *ProteoSeeker* includes a custom method of taxonomy assignment based on screening each protein against the TPD and in case of at least one hit, screening the protein in turn against the TFPD and identifying the best hit and the taxa associated with it. Then, each bin, based on its proteins and their taxonomy classifications, is associated with one or more taxa, and then all genes and proteins of that bin are assigned the same taxa. The nr database was used as the reference protein database for filtering. In addition, initial testing was performed on using the Uniref50 and Uniref90 as the protein databases for filtering, as they are of significantly smaller sizes. Due to the reason that many headers of proteins from these Uniref databases include general taxonomy information (e.g., “root”, “bacteria”), the results were not as encouraging as the results acquired by utilizing nr as the protein database to be filtered. *ProteoSeeker*, however, is designed to also handle and extract the taxonomy information provided in the headers of proteins based on the header-style of the Uniref databases.

One part for consideration regarding the seek mode of *ProteoSeeker* is that it should be able to adequately recognize each protein family represented by a single Pfam profile, unique to the family. Selecting a protein family with at least one domain not unique to the family, will lead *ProteoSeeker* searching for all families associated with that domain. While a protein belonging to such types of protein families should be identified by *ProteoSeeker,* it could be mixed with other proteins each comprising a different set of domains including at least one associated with the selected protein families. Despite this, the additional information provided by *ProteoSeeker* facilitates the process of discarding certain proteins. For example, such a helpful step in the latter process is the evaluation of the protein length of each of the identified proteins compared to the protein family of its best hit against the UniprotKB/Swiss-Prot database and the provided mean length of the latter family. In addition, the presence or absence of signal peptides and transmembrane domains is another piece of information that might help the user decide whether a predicted protein belongs to the targeted protein family. Furthermore, a protein discovered by *ProteoSeeker*, successfully passing the stage of the profile screening, will contain at least one of the domains corresponding to the selected protein families but not necessarily all. Protein families, which are not represented by any type of profile in Pfam, remain a blind spot in the screening search area of *ProteoSeeker* while running in seek mode of type 1 analysis.

The usefulness of *ProteoSeeker* and its effectiveness in identifying proteins carrying targeted functionalities has been already demonstrated by the discovery of novel amylase and CA enzymes with predefined biochemical characteristics. Interestingly, in the case of the CA-KR1 discovery, the novel biocatalyst consists of a high-profile enzyme with great potential for carbon capture applications, which has already received significant attention by the relevant community.

The taxonomy route of COMEBin/MetaBinner of *ProteoSeeker* was created as a combination of processes included at the already existing by then seek mode and binning methods of COMEBin and MetaBinner. The evaluation of the taxonomy mode of *ProteoSeeker* was based on artificially created (simulated) reads and on reads originating from known mixtures of microorganisms (cultures), with the addition or not of specific biases in certain cases. The protein database selected to be used in the taxonomy route of COMEBin/MetaBinner was the nr database. Other elements were considered in the evaluation, such as the database used in the route of Kraken2 as well as different filtering thresholds for the post-filtering analysis of the species identified by Kraken2 and processed by Bracken. The results of the evaluation offered a series of interesting observations.

The evaluation results were studied according to the scores of the metrics for the different routes, tools, databases and filtering thresholds and in regards of the biases and origin of the samples (Figure 4 and Additional file 1: Figures S2 and S3). An initial observation is that the scores of the same combinations of taxonomy routes, tools, databases and filtering thresholds seem to follow the same pattern between the different samples of each species-abundance category. This pattern seems to be most disrupted for the two groups of low-species numbers, meaning 10 and 40, and to be most evident for the groups containing high numbers of species and especially for the “control” samples which originate from cultures. This is an expected outcome since samples containing a few species, based on a few differences regarding their true positive, false positive and false negative hits, will show large discrepancies between their scores regarding the rest of the metrics. The same applies for the L1 norm metric, which also accounts for the relative abundances of the species. In addition, the control samples can be grouped in two categories, each containing two samples, each of which originates from NGS analysis of the same library preparation. The validation group of species and their relative abundances for each of the control samples is the same and therefore the results of the taxonomy routes were expected to be the most similar for these samples. In addition, most combinations of Kraken2 databases and filtering thresholds, for most samples, showed increased sensitivity, precision and accuracy for the GC-rich samples compared to the same samples with no bias and with AT-rich bias. The results from the COMEBin/MetaBinner route did not reveal any evident pattern regarding the biases of the samples.

Furthermore, based on the best scoring combinations of Kraken2 databases and filtering thresholds, a highly significant result is that the Kraken2 Standard-8 and Standard-16 collections can in general provide adequate and, in specific cases, the best results when combined with the right filtering threshold, even compared to the Standard collection. Several combinations of these databases with different filtering thresholds were present in the top 2-3 combinations for different metrics, as shown in Additional file 2. This is a crucial observation because the Standard-8 and Standard-16 collections do not require large amounts of memory space as the Standard collection or other databases demand.

The selected best scoring Kraken2 combinations, based on each metric, involve the database of the Standard-8 collection with the filtering threshold of 5.0% and the filtering threshold computed for non-gut samples, the database of the Standard-16 collection with the filtering threshold of 5.0% and the database of the Standard collection without a filtering threshold and with the filtering thresholds of 100 and 5.0%. Of all combinations, the Standard-8 collection combined with the filtering threshold computed for non-gut samples is for most samples in the top 2 or 3 scoring methods offering a consistent performance across all metrics. The same combination has the best performance for accuracy for most samples. The Standard-8 collection with the filtering threshold computed for non-gut samples and the Standard collection without a filtering threshold and with the filtering threshold of 100 have the best performance for sensitivity. The Standard-8, Standard-16 and Standard collections, each with the filtering threshold of 5.0%, have the best performances for most samples for precision. Furthermore, the combinations of the Standard-8 collection with the filtering threshold computed for non-gut samples and the Standard collection with the filtering threshold of 100 are in the top 2-3 scoring combinations for most samples based on the F1 score, Jaccard index and L1 norm. The latter metric is the one that also accounts for the relative abundances of the species.

While computing the different metrics, the species taken into account by the taxonomy route of COMEBin/MetaBinner come from the bins associated with one species each. The relative abundance of each of these species is the sum of the relative abundances of the same species based on all bins that were associated with that species alone. The taxonomy route of COMEBin/MetaBinner by analyzing solely its own performance, scored in general low for sensitivity and accuracy and much higher for precision. Based on the F1 score and Jaccard index, COMEBin and MetaBinner scored moderately and based on the L1 norm they scored poorly relative to the Kraken2 taxonomy route.

In general, it should be noted that the combinations of the Kraken2 databases of the Standard-8, Standard-16 and Standard collections with the 5.0% filtering threshold seem to have very close scores in all metrics and samples. Based on closer examination of the results, the number of species predicted by Kraken2 in each sample is increased as the database becomes larger in size, but the species remaining after the application of the filtering threshold are for almost all samples the same and with approximately the same relative abundances. Therefore, in this case, it appears that most of the species predicted with low relative abundances by larger databases are discarded with a relatively high filtering threshold, forming a set of species more similar to the ones predicted by smaller databases.

Hence, the taxonomy route of Kraken2 with the proper filtering threshold performs better in terms of species identification and quantification for a sample compared to the route of COMEBin/MetaBinner. There are two main reasons for the latter route being a necessary part of the pipeline: the first reason is that the process of binning is a crucial analysis process of analyzing a metagenomic dataset regardless of whether the taxonomy takes place after or not. The taxonomy process itself, as shown by the stage-specific execution times of *ProteoSeeker*, is a much faster process than binning and does not act as a bottleneck to the pipeline, even though it could be omitted completely. We believe that for metagenomic datasets including a multitude of microorganisms, whose genomes are yet to be identified and documented, binning the reads may be a more appropriate approach than trying to directly assign species to them primarily based on information originating from known and documented genomes. Binning is in part based on general biological factors, which are used to group contigs coming from the same organisms without including comparisons with known sequences. Hence, each bin may later act as groups of contigs, in turn of reads, genes and proteins originating from the same organism. The second reason is that the taxonomy route of COMEBin/MetaBinner appears to be efficient in the taxonomy classification of proteins from known protein families in a sample as it can base the taxonomy process on protein families associated with the ones which the proteins of interest are expected to belong to. More specifically, for each enzyme tested with its functionally confirmed experimentally, originating from a sample with a FASTQ dataset available, the taxonomy classification of its best hit (with the lowest E-value) from the results obtained by screening it against the nr database of NCBI through the online blastp suite of NCBI was acquired. The latter classification was set as the correct classification for each enzyme and it was compared to the classification provided for the same enzyme by both taxonomy routes of *ProteoSeeker*. It should be emphasized that the COMEBin/MetaBinner route is more biased towards making a correct classification compared to the Kraken2 route, because the COMEBin/MetaBinner route utilizes a filtered protein database formed based on the nr database which was also used by the blastp suite, while the databases used by the Kraken2 route are the Kraken 2 / Bracken Refseq indexes of the Standard-8, Standard-16 and Standard collections. The taxonomy route of COMEBin/MetaBinner, for each enzyme and both COMEBin and MetaBinner, except for the case of MetaBinner for CA-KR1, was able to identify the correct species or genus. According to the results of the Kraken2 taxonomy route, the taxonomy classification of CA-KR1 was not inferred based on the databases of the Standard-8 and Standard-16 collections and the species and genus predicted based on the database of the Standard collection did not match the target species and genus, respectively. For CA_201 and AL_17, an incorrect species of the right genus was identified for each case of database. It should be noted that the relative abundances of the species associated with the CA-KR1, CA_201 and AL_17 enzymes by the Kraken2 route are all below 0.1%, while by the COMEBin/MetaBinner route are above 0.1%. The reason for the COMEBin/MetaBinner route to misclassify the enzymes while basing its analysis on a filtered nr database, could be based on the protein of the best-hit of an enzyme against the non-filtered nr, being omitted during the filtering process, or the protein having no hits against the TPD or the classification of the bin which contains the enzyme being different than the correct one due to another classification having the highest frequency for that bin based on its proteins.

In general, we highly recommend using the Kraken2 taxonomy route to perform the taxonomy classification of the proteins identified in a sample, while it is also worth performing solely or additionally the COMEBin/MetaBinner taxonomy route based on the nr database when interested in the taxonomy classification of proteins, whose protein families are well known/studied or/and multiple proteins of these protein families exist in the nr database. When running *ProteoSeeker* based on the taxonomy mode and COMEBin/MetaBinner route, providing protein family codes, for families which are associated with the families in which the proteins of interest are expected to be part of, increases the chances of performing a successful or at least promising taxonomy prediction for the proteins of interest when applying the COMEBin/MetaBinner route. We also recommend using the Kraken2 taxonomy route with any of the Standard collections by providing multiple filtering thresholds and selecting to bin the reads based on the filtering threshold computed for non-gut or gut samples. Examining the filtered species based on the different thresholds provided, in general provides an insight to the user about which threshold is the most suitable one based on the sample being analyzed and using that threshold to re-bin the contigs in another run. Binning the contigs and in turn reads, can easily be performed later again based on a new filtering threshold by omitting the processes prior to the binning stage.

As showcased in this study, regarding the process of taxonomy classification, the user is able to prioritize sensitivity, accuracy, precision, F1 score, the Jaccard index or L1 norm or identifying the protein families of proteins expected to belong to specific protein families, by applying either the Kraken2 taxonomy route based on a database and a filtering threshold or the COMEBin/MetaBinner taxonomy route based on a protein database, a set of protein families, a set of protein names and a minimum length for the contigs to be binned. In principle, the COMEBin/MetaBinner taxonomy route scored moderately in relation to the selected best scoring Kraken2 combinations of databases and filtering thresholds, for most metrics and samples.

Utilizing the nr database in the taxonomy route of COMEBin/MetaBinner is a difficult task from the prospect of space availability. As part of future work, we are planning to reform the Uniref50 and Uniref90 databases, by including in the headers of their proteins the species documented for those proteins based on the information available in the UniProtKB/Swiss-Prot protein database. These reformed protein databases could be tested on whether they could be used as alternative options to the nr database for the taxonomy route of COMEBin/MetaBinner of the taxonomy mode or the type 2 analysis of the seek mode of *ProteoSeeker*.

Total and stage-specific execution times for ProteoSeeker were computed based on the runs of analyzing the 19 gold standard samples for its taxonomy evaluation. The total execution times of *ProteoSeeker* do not include the time spent for downloading and converting the SRA dataset to FASTQ and filtering the protein database for the COMEBin/MetaBinner route. In addition, the total execution time of *ProteoSeeker* for the species-abundance categories of 10, 40, 120, 500 and 1000 is at most 61.13 min, 43 min, 120.63 min, 141.44 min and 312.73 min respectively. There is evident variety regarding the total execution time of *ProteoSeeker* between the different samples of each category. This variety cannot be explained based on the size of the FASTQ files analyzed by *ProteoSeeker* for each sample. It can be attributed mainly on the stage-specific execution times of the assembly and binning stages. Based on the stage-specific execution times of *ProteoSeeker* the assembly and binning stages can be identified as the most time-consuming steps and main portions of the total execution times, except for the taxonomy analysis in the samples of 10 species where the taxonomy stage occupies a significant part of the total execution time while remaining below 10 min. It should be noted that the stages prior to and including the gene prediction stage, are common between the different taxonomy methods of each sample as they were performed once, only for the run based on the Kraken2 route with the database of the Standard-8 collection. In general, we can observe that the time needed for the assembly increases as the number of species in the samples also increases. The binning stage seems to demand approximately the same portion of time as the assembly for the COMEBin/MetaBinner route. For the Kraken2 route, binning is performed much faster. For the COMEBin/MetaBinner route, in our case, binning depends on the number of contigs surpassing the threshold of 1,000 nucleotides. It seems that the higher the number of the latter contigs, the higher the binning time, regardless of the initial size of the sample. In general, for each sample the execution time of *ProteoSeeker* is increased from the smallest Kraken2 database to the largest, then to MetaBinner and then to COMEBin. One should of course take into consideration the fact that Kraken2 was run without memory mapping and COMEBin was run by utilizing a GPU. In the opposite case for either tool, running Kraken2 with memory mapping or COMEBin without a GPU significantly increases the time of the taxonomy stage and the time for the binning stage respectively.

The analysis based on the taxonomy evaluation for the total execution time of *ProteoSeeker* in relation with the size and the species-abundance of the samples, evidently showed that the execution time is proportionally dependent on the species-abundance of the sample. It should be noted that that the execution time is proportional to the species-abundance of the samples including only the samples of simulated origin.

Time execution analysis was not performed solely for the seek mode of *ProteoSeeker*. The time execution analysis for the taxonomy mode of *ProteoSeeker* was a more valid approach due to the existence of the gold standard samples with the simulated reads corresponded to specific species with known relative abundances and biases, which allowed for a controlled analysis to take place. The latter analysis refers to the ability of making connections between the total and stage-specific execution times of *ProteoSeeker* with the biases as well as the sizes and species-abundances of the samples. In addition, most of the stages of the seek mode are identical with those of the taxonomy mode, e.g., SRA to FASTQ conversion, creation of a profile and a filtered protein database, quality control, preprocessing the reads, assembly, gene prediction and annotation, protein clustering. Therefore, any observations made regarding the time execution analysis of these stages based on the taxonomy mode and its evaluation are also applied to the same stages of the seek mode. Furthermore, we are noting certain observations we have made throughout the application of *ProteoSeeker* in numerous datasets up to this day, including its runs on datasets related to the enzyme discovery projects. These observations are related to both the seek and taxonomy modes of *ProteoSeeker* and focus mainly on the later stages of the seek mode for which no observations were made through the taxonomy evaluation. Providing as input numerous protein families or generic protein names to *ProteoSeeker* may lead to the creation of large SPD, TPD, SFPD and TFPD databases, which will cost additional time regarding the stages that include their screening in both the seek and taxonomy modes. Abbreviations used as protein names should be encompassed by empty spaces (e.g., to use “CA” as the abbreviation for CAs as a protein name it should be provided as “ CA “). Screening a large SPD or TPD database is much less time-consuming that screening a large SFPD or TFPD database. The seek mode of type 1 analysis is based on screening the SPD, while the type 2 analysis is based on screening the SFPD. In addition, the taxonomy route of Kraken2 is not affected by the sizes of these databases but the taxonomy route of COMEBin/MetaBinner is, specifically by the TPD and TFPD databases. Screening the Pfam database in the seek mode is relatively fast except if the number of proteins reaching that stage is exceptionally large. Using CD-HIT to reduce that number by clustering the proteins, without losing valuable information, is both easy to apply and time-efficient, as ProteoSeeker can directly be applied after the gene prediction stage (where gene annotation and applying CD-HIT follow) and also CD-HIT is an algorithm whose application is in general fast. In addition, most often, the protein family identification according to the best hits of the proteins based on their screening against the UniprotKB/Swiss-Prot database, the screening for input motifs and the topology predictions made by Phobius are rapid processes, which we have never encountered as bottlenecks for *ProteoSeeker*.

## Conclusions

In response to the lack of standardized guidelines, the complexity and general demand in computational resources of existing pipelines, in the field of metagenomic analysis of WGS data, we have developed *ProteoSeeker*, a rapid and precise analytical tool for these purposes. In addition, ProteoSeeker can process genomic (genomes or contigs) and proteomic data (protein sequences in FASTA format). Intended also for application by non-expert users, *ProteoSeeker* integrates state-of-the-art tools and automates workflows based on minimal user input. Surpassing the ability of each integrated tool, *ProteoSeeker* enhances accessibility and consistency across studies with the scope of identifying proteins of interest. *ProteoSeeker* allows the user to easily modify the behavior or even the type of the tools and the databases utilized in its pipeline. *ProteoSeeker* screens a dataset and identifies proteins that belong to user-defined protein families while performing taxonomic analysis. *ProteoSeeker* has been already utilized successfully to discover novel enzymes for industrial applications with *a priori* desirable functionalities, including a highly promising CA biocatalyst for biomimetic carbon capture (36). The latter analyses and discoveries support the usefulness of *ProteoSeeker’s* seek mode and, in turn, of its analysis types that include capability to be molded based on user-input information leading to protein discovery and annotation regarding specific protein families.

Regarding the taxonomy mode of ProteoSeeker and based on the evaluations of the same mode, in general, the Kraken2 taxonomy route is highly advised to perform the taxonomy classification, while the COMEBin/MetaBinner taxonomy route can additionally provide valuable information when searching for proteins of specific protein families while providing those families, as well as families or/and protein names associated with them as input to *ProteoSeeker*. The user can focus on any of the metrics used for the evaluations (true positive hits, false positive hits, false negative hits, sensitivity, accuracy, precision, F1 score, Jaccard index, L1 norm) by applying the Kraken2 taxonomy route based on a database and a series of filtering thresholds. The examination of the results provided for each filtering threshold can lead the user to identify the one most suitable for the analysis and easily configure and re-run *ProteoSeeker* by basing binning and subsequent stages of the taxonomy mode on the selected filtering threshold.

The total execution times computed for *ProteoSeeker* based on its taxonomy evaluation showed that the assembly and binning stages can be identified as the most time- consuming steps and main portions of the total execution times for its taxonomy mode. The binning stage for the COMEBin/MetaBinner route can be affected by the specified minimum length of the contigs to be used for binning. Most initial stages of the seek mode are identical with the respective ones from the taxonomy mode, thus our observations relating the total and stage-specific execution times of *ProteoSeeker* to the gold standard samples, are common between both modes for those stages. Selecting numerous protein families or protein names which may be associated with multiple protein families may lead to the creation of large SPD, TPD, SFPD and TFPD databases, potentially as large at the protein database to be filtered itself. For that reason, it is important, when in need of selecting high number of protein families or protein names, to configure *ProteoSeeker* to stop after creating these databases and examine the latter regarding their size or/and contents. Then one can re-run *ProteoSeeker* after the stage of creating the profile and filtered protein databases. The size of the SPD and TPD will affect the execution time of their screening in the type 1 analysis of the seek mode and in the taxonomy part of the COMEBin/MetaBinner route of the taxonomy mode, respectively. The size of the SFPD and TFPD will affect the execution time of their screening in the type 2 analysis of the seek mode and in the taxonomy part of the COMEBin/MetaBinner route of the taxonomy mode. CD-HIT is a crucial stage of the pipeline regarding controlling the number of proteins which are used in the subsequent stages by utilizing it to cluster the proteins and retaining the representatives of the clusters. Reducing the redundancy of the protein sequences without losing valuable information from the sample is additionally a method for decreasing the execution time of the screenings of the SPD, TPD, SFPD and TFPD databases.

With the introduction of *ProteoSeeker* to the biotechnology community, we anticipate a transformative impact on metagenomic research. This comprehensive tool is designed to accelerate the discovery of novel biocatalysts and biomolecules, leveraging the ever-growing repository of publicly available metagenomic data to drive biotechnological innovation forward. Emphasizing accessibility and automation, *ProteoSeeker* is positioned to become an indispensable tool for bioscientists and biotechnologists working on protein discovery. Widespread adoption of *ProteoSeeker* is poised to accelerate the discovery of novel enzymes and biomolecules, propelling innovation in biotechnological research across multiple sectors.

## Methods

*ProteoSeeker* implements a new pipeline targeted at the analysis of metagenomic data from WGS and by extension of genomic (genomes or contigs) and proteomic data. The pipeline incorporates multiple bioinformatic analysis tools, offering their automated synergy and providing multiple options for its versatile application. The functions of *ProteoSeeker* are dependent on the existence of certain datasets. These datasets have been the result of performing an analysis on the UniprotKB/Swiss-Prot protein database. Initially, the reviewed proteins of the UniprotKB/Swiss-Prot database were downloaded and the length, the organism, the name(s), the Pfam profiles and the protein family assigned to each protein were collected. Each protein family was associated with a group of proteins. Each protein was associated with a set of Pfam profiles where each profile has a frequency. Two sets of profiles are considered to be identical if their profiles and their frequencies match. The frequency for each set of profiles of the proteins in a protein family was computed. Consequently, each protein family, through its protein group, was associated with one or more sets of profiles of maximum frequency within the protein group, a mean and median length, and a set of protein names. Each protein name was also accompanied by a frequency. Two sets of profiles, from two different proteins, are identical when each profile of one set is of the same type and has the same frequency as another profile of the other set and vice versa. The different sets of profiles which may have been associated with a protein family at this point may include profiles of the same type.

*ProteoSeeker* offers two primary and independent functionalities. The first functionality is termed “seek functionality” and it is applied through the “seek mode” of *ProteoSeeker* (Figure 2). This mode includes three types of analysis, termed “type 1”, “type 2” and “type 3” respectively. Type 1 analysis includes searching for putative proteins which contain domains corresponding to the profiles associated with one of the selected protein families. Type 2 analysis includes searching for putative proteins which contain none of the domains used for screening in type 1 analysis and have at least one hit of low enough E-value against the “seek filtered protein database” (SFPD). Type 3 analysis includes both type 1 and type 2 analysis. The second functionality is termed “taxonomy functionality” and it is applied through the “taxonomy mode” of *ProteoSeeker* (Figure 3). It includes two distinct taxonomy routes each of which performs a taxonomy analysis of the putative proteins and binning of the contigs. The first route, the “Kraken2 route”, performs taxonomy classification of the reads through Kraken2, abundance estimation through Bracken and then bins the contigs based on the taxonomy classification of their reads. The second route, the “COMEBin/MetaBinner route”, at first bins the contigs, then isolates the proteins which include at least one domain corresponding to a profile from the TPD, runs DIAMOND to screen the latter proteins against the TFPD, determines the taxon or taxa associated with the best hit of each protein and eventually assigns one or more taxa to a bin and its genes and proteins.

All stages of the pipeline implemented by *ProteoSeeker* will be described based on the application of its processes for a type 3 analysis (both type 1 and type 2 analyses) of the seek mode and both taxonomy routes of the taxonomy mode.

Based on the datasets provided from the analysis of the UniprotKB/Swiss-Prot database, each protein family selected corresponds to one or more sets of Pfam profiles based on which in turn a unique set of profiles is formed. The latter set is used by HMMER to create a database of profiles (48,49). Selecting protein families for the seek or taxonomy mode leads to the creation of a “seek profile database” (SPD) or a “taxonomy profile database” (TPD) respectively. *ProteoSeeker* may also accept as input a profile database directly. When selecting protein families, the following stage is to utilize a protein database. The protein database selected as the most suitable one for the analyses performed by ProteoSeeker is the non-redundant (nr) database of NCBI. The protein database is filtered based on the protein names associated with the selected protein families. The protein names of each selected protein family are filtered based on a threshold. This threshold is a percentage and computes the value of the minimum frequency required for a protein name not to be discarded, based on the maximum frequency of the protein names. Each protein name that surpasses the threshold is used in filtering the protein database and creating either the “seek filtered protein database” (SFPD) or the “taxonomy filtered protein database” (TFPD). This filtering is carried out by a supportive stand-alone Python command-line tool and module, which can take as input sets of protein names and corresponding file names and can run on multiple processes in parallel. The SFPD or TFPD is converted to a database by DIAMOND. The user may provide a protein file and its corresponding database directly. Also, the user can provide his own set of protein names and select whether only the names he provided will be used to create the SFPD or TFPD, or the names he provided will be added (if not present already) to the ones associated with the selected protein families. Utilizing a protein database is mandatory when the type 2 analysis of the seek mode is to be performed or when taxonomy is to be carried out through the taxonomy route of COMEBin/MetaBinner. All processes related to utilizing the selected protein families for the taxonomy mode (e.g., creating the TPD, filtering the protein database and creating the TFPD) are omitted when the taxonomic analysis is based on the taxonomy route of Kraken2. Using Kraken2 comes with the need for indexes which come in a variety of composition and size and can either be downloaded as pre-built databases or be user-built databases.

One mandatory input for *ProteoSeeker* is an SRA code (RUN accession) from the SRA database of NCBI or a dataset (file input). If an SRA code is provided, then *ProteoSeeker* automatically downloads the SRA file associated with the code, validates its integrity, and converts it to its corresponding compressed FASTQ file(s). In the case of file input, this may come as one FASTQ file with single-end reads from Next-Generation Sequencing (NGS) or as two FASTQ files with paired-end reads from NGS or as a FASTA file containing already assembled reads (contigs) or genomes or as a FASTA file containing protein sequences. Based on each case of file input, specific early stages of the pipeline implemented in *ProteoSeeker* are omitted. Hence, *ProteoSeeker* can actually process metagenomic data from WGS, genomic and proteomic datasets.

Both modes of *ProteoSeeker* begin with running FastQC (50) to perform several quality control checks of the reads. A brief analysis of the results from FastQC follows which is targeted on the number of the reads and the overrepresented sequences identified. The next stage of the analysis is preprocessing the reads with BBDuk (51) which involves filtering the reads based on quality and length examinations and trimming the reads based on a list of sequences (“adapters file”). A user can provide a file which contains adapter sequences or use an existing one which contains adapters generally used by various NGS platforms. The user can also select whether the overrepresented sequences identified by FastQC will be added in the adapters file. Assembly is the next crucial stage of *ProteoSeeker*. It is performed by MEGAHIT (52) and provides contigs as part of the output.

The following processes are solely part of the taxonomy mode. The taxonomy route of COMEBin/MetaBinner at this stage uses the binning tool of COMEBin or MetaBinner to bin the contigs. Binning is based on the contigs and the preprocessed FASTQ files. The taxonomy route of Kraken2 at this stage uses Kraken2 which based on a Kraken2 database predicts species by analyzing the reads of the sample and assigns these species to the reads. The abundances and relative abundances of the Kraken2 predicted species, at the species level, are estimated by Bracken. The same abundances are filtered based on one or more filtering thresholds, if any provided. A filtering threshold may target the abundances or relative abundance of the species. Any species not surpassing a threshold is discarded. In this filtering process *ProteoSeeker* is also able to determine automatically the filtering threshold for “non-gut” or “gut” samples, according to the Shannon index computed based on the results of Bracken, as described by the method and formula proposed by Poussin *et al.* (40). Each filtering threshold applied leads to a filtered list of species. One of the filtering thresholds (can be user-defined) is used by *ProteoSeeker* to select the corresponding set of filtered species to be used by certain subsequent stages of the analysis (e.g., binning). The Shannon index is computed based on the KrakenTools.

The next step of *ProteoSeeker*, for both its modes, is the application of the gene prediction tool, FragGeneScanRs (53). This tool can identify protein coding regions, including prokaryotic genes, and provide their protein sequences. Moreover, custom translation tables based on NCBI’s genetic codes are provided for usage. Information is also collected for each gene regarding the presence or not of start and end codons, its length, and its distance from the ends of its contig. CD-HIT (54,55) is then applied to perform clustering of the protein sequences. It can handle extremely large databases and can help to reduce the computational demands of subsequent stages by reducing the redundancy of the proteins.

The following steps are part solely of the seek mode of *ProteoSeeker*. *ProteoSeeker* proceeds, in type 1 analysis, into screening the proteins against the SPD with HMMER. The proteins subject for further annotation, contain each protein that scored at least one hit against the SPD (protein set 1). Additionally, in type 2 analysis, the rest of the proteins (not part of protein set 1) are screened against the SFPD by DIAMOND. Those that scored at least one hit with E-value equal to or lower than a specific threshold (by default 1e-70), are also subject for further annotation (protein set 2). Both protein sets are then combined and subjected to screening, through HMMER, against all the profiles of the Pfam database. Furthermore, protein set 1 (if any) is screened against the SFPD by DIAMOND. Both protein sets are also screened by DIAMOND against the UniProtKB/Swiss-Prot protein database.

The following steps are part solely of the taxonomy mode of *ProteoSeeker*. At this point, a stage common between both taxonomy routes is that of read mapping. Reads are mapped to contigs through Bowtie2 (56,57). For the COMEBin/MetaBinner taxonomy route each bin and its taxa are both quantified based on the reads mapped to the contigs of the bin. The relative abundance of a bin and of its taxa is the proportion of the reads mapped to the contigs of the bin against the total number of preprocessed reads. For the taxonomy route of COMEBin/MetaBinner *ProteoSeeker* performs screening of the proteins against the TPD. Proteins with at least one hit are then screened through DIAMOND against the TFPD. The TFPD database is parsed, and every protein is associated with one or more taxa based on the information in its header. In turn, each putative protein is associated with one or more taxa based on its hit (if any) of the lowest E-value against the TFPD. The TaxIds of the latter taxa and their lineages are found based on TaxonKit (58). Each contig contains one or more genes, thus is associated with their proteins. In turn, each bin contains several contigs and is associated with their proteins. By using proteins who were assigned at least one taxon and based on the proteins included in a bin, each bin is assigned one or more taxa accompanied by frequencies. The frequency of each taxon is the number of times this taxon is found associated with the proteins of the bin. The taxon or taxa with the highest frequency for the bin are identified and are assigned to that bin. The same taxon or taxa are also assigned to every gene and protein of the bin. In the Kraken2 taxonomy route, by combining the information from the filtered (non-discarded) species assigned to the reads and the read mapping to the contigs by Bowtie2, species are associated with the contigs. Each species associated with a contig is accompanied by a frequency, equal to the number of reads associated with that species and mapped to that contig. If a single species for a contig has the highest frequency, then that species is assigned to the contig, otherwise no species is assigned to the contig. Based on the species of the contigs, the latter are binned. Hence, each bin of contigs is directly assigned a single species and also quantified based on the quantification provided initially by Kraken2 for the same species. The next step is the assignment of the species of a bin to each gene and protein of that bin.

In the case of applying solely the taxonomy mode of *ProteoSeeker* the pipeline does not perform any other analysis after this stage. Otherwise, the pipeline goes on to perform more analyses in the seek mode. The first analysis is to predict the transmembrane topology of the proteins of both protein sets, by Phobius (59). Phobius is a tool that performs transmembrane topology and signal peptide prediction in proteins. The second analysis is to search for user-input motifs in the protein. The third analysis is to predict the protein family of each putative protein. This is accomplished by determining the hit of lowest E-value acquired by screening the putative protein against the UniprotKB/Swiss- Prot database through DIAMOND and identifying the protein family of that hit. The mean and median length of the latter family is determined based on the datasets provided from the initial analysis of the UniprotKB/Swiss-Prot database. Moreover, the length of the putative protein is compared with the mean length of the predicted protein family and their difference is documented in the results and final annotation.

Lastly, for both the seek and taxonomy modes, annotation files are generated by *ProteoSeeker*. These files contain a synopsis of the information collected for each putative protein from either or both of the two modes. A summary of the type of results found in the annotation files can be found in Additional file 3. The user can specify which information is to be included in the annotation files based on the seek mode or the taxonomy mode or both modes. In addition, if *ProteoSeeker* is run after or up to a specific stage of the pipeline for a sample or dataset already analyzed previously, *ProteoSeeker* collects available information from the previous run(s), combines them and outputs the previous and new information in a new set of annotation files. This is possible as at each stage of the pipeline *ProteoSeeker* stores the information collected for the putative proteins in specific files which it analyzes in future runs of the same sample or dataset. It should be noted that, at each possible termination point of *ProteoSeeker*, the information for the execution times of the different stages of *ProteoSeeker* which have been run in the pipeline are provided, as well as for the whole run.

### Gold standard datasets used in the evaluation of the “taxonomy” mode of *ProteoSeeker*

In total, 19 SRA files corresponding to synthetic datasets or datasets from the NGS of samples from cultures of known composition of species were used in the evaluation of the taxonomy mode of *ProteoSeeker* and the analysis of its execution time. More specifically, 15 samples have been generated *in silico* simulating NGS reads from an Illumina HiSeq4000 sequencer (2 x 150-bp paired-end reads). In each of the latter samples a number of reads has also been added from sequencing mouse cecal samples. Furthermore, certain samples were formed with the bias of being AT-rich or GC-rich. The remaining 4 samples of the total 19 samples, originated from the commercially available ZyMoBIOMICS^™ Microbial Community DNA Standards and were sequenced in Illumina HiSeq4000 generating 2 x 151-bp paired-end reads. The creation process of the 15 artificial samples and the sequencing process of the other 4 samples were not conducted in our study. The latter processes are described in detail in the work of Poussin *et al.* (40).

## Abbreviations

WGS: Whole-Genome Sequencing
CA: Carbonic Anhydrase
CPU: Central Processing Unit
GPU: Graphics Processing Unit
NGS: Next-Generation Sequencing
SPD: seek profile database
TPD: taxonomy profile database
SFPD: seek filtered protein database
TFPD: taxonomy filtered protein database
HMM: hidden Markov model

## Availability of data and materials

The datasets, source code of version 1.0.0 of ProteoSeeker, the versions of the tools included in the *ProteoSeeker* pipeline of version 1.0.0 and the collection dates of the databases which support the conclusions of this article are publicly available in the GitHub repository of *ProteoSeeker* with the tag “v1.0.0” at “https://github.com/SkretasLab/ProteoSeeker” (25) and with the version 1.0.0 used in the manuscript deposited in the DOI-assigning repository Zenodo at “https://doi.org/10.5281/zenodo.13944968” (60). *ProteoSeeker* is also shipped as a docker image through Docker Hub by the repository “skretaslab/proteoseeker” at “https://hub.docker.com/r/skretaslab/proteoseeker” (26). To access the website of *ProteoSeeker* please visit “www.skretaslab.gr/proteoseeker” (27). The datasets of the gold standard samples used in the evaluation are described in the work of Poussin *et al.* (40), their SRA accession numbers and other related information are provided in Additional file 1: Table S1.

Project name: ProteoSeeker Project home page: https://github.com/SkretasLab/ProteoSeeker (25) and www.skretaslab.gr/proteoseeker (27) Archived version: https://doi.org/10.5281/zenodo.13944968 (60) Operating system(s): The Linux operating system is needed to install and run the command-line tool. The Linux operating system with the amd64 architecture and a platform that supports running Docker images is needed to run the Docker image.

Programming language: Python, Bash

Other requirements: The Anaconda environment is needed for the command-line tool of ProteoSeeker. A system capable of deploying Docker containers from Docker images is needed for the Docker image of ProteoSeeker.

License: GNU General Public License (GPLv3) for the source code of ProteoSeeker.

## Acknowledgements

None.

## Author information

Authors and Affiliations

Institute for Bio-innovation, Biomedical Sciences Research Centre “Alexander, Fleming”,

Vari, Greece

Georgios Filis, Dimitra Bezantakou, Konstantinos Rigkos, Dimitra Zarafeta & Georgios Skretas

Institute of Chemical Biology, National Hellenic Research Foundation, Athens, Greece

Georgios Filis, Konstantinos Rigkos, Despina Noti, Pavlos Saridis, Dimitra Zarafeta & Georgios Skretas

Department of Informatics and Telecommunications, National and Kapodistrian University of Athens, Athens, Greece

Georgios Filis

Department of Biological Applications and Technologies, University of Ioannina, Ioannina, Greece

Konstantinos Rigkos

Faculty of Biology, National and Kapodistrian University of Athens, Athens Greece

Pavlos Saridis

Contributions

GS and DZ designed and managed the project. GF wrote the source code of ProteoSeeker, and the code used for the evaluations of ProteoSeeker. GF and DB wrote the documentation for ProteoSeeker. GF and DB performed the evaluations of the seek and taxonomy modes of ProteoSeeker. KR performed experimental analyses of CAs and ALs. DN performed experimental analyses of ALs. PS performed experimental analyses of CAs. GF, DB, KR, DN, DZ, GS wrote the manuscript. DZ and GS revised the manuscript. All authors wrote and approved the final manuscript.

Corresponding authors

Correspondence to Dimitra Zarafeta and/or Georgios Skretas.

Ethics declarations

Ethics approval and consent to participate

Not applicable.

Consent for publication

Not applicable.

Competing interests

The authors declare that they have no competing interests.

## Supplementary Information

Additional file 1

PDF

ProteoSeeker: A feature-rich metagenomic analysis tool for Accessible and Comprehensive Metagenomic Exploration. Supplementary Text, Supplementary Figures and Supplementary Tables.

Contains a description of the filtering process of a protein database which may be performed during a run of ProteoSeeker. Contains a description of the metrics used in the evaluation of the taxonomy mode (taxonomy routes) of ProteoSeeker. It also contains the following figures and tables:

Fig. S1. Biochemical characterization of IMAC purified AL_6 and AL_15. Fig. S2. True positive, false positive and false negative hits of the taxonomy mode evaluation results with the samples sorted based on their species-abundances and biases. Fig. S3. F1 score, Jaccard index and L1 norm of the taxonomy mode evaluation results with the samples sorted based on their species-abundances and biases. Fig. S4. The total execution time of *ProteoSeeker*, based on its taxonomy mode evaluation, for each database and in turn for each of the 19 gold standard datasets, relative to their sizes. Fig. S5. The mean execution time of *ProteoSeeker*, based on its taxonomy mode evaluation, for each database and in turn for each of the 19 gold standard datasets, relative to their species abundance. Table S1. Information about each of the 19 gold standard datasets. Table S2. Information about the different evaluation cases of the taxonomy mode (or taxonomy routes) of *ProteoSeeker*. Table S3. Information for the taxonomy classification of experimentally verified and studied enzymes CA-KR1, CA_201 and AL_17, discovered by the seek mode of *ProteoSeeker*.

Additional file 2 TSV

**The frequency of best scores of each combination of Kraken2 database and filtering threshold used in the taxonomy evaluation for each metric across all 19 samples.**

Contains for each metric and each taxonomy route, every combination of taxonomy tool and database used in the run, and a frequency for that combination which is based on the number of times that combination had the best score for the metric according to each of the 19 gold standard samples.

**Additional file 3**

**TSV**

**A summary of the results documented in the annotation files generated from a**

***ProteoSeeker* run after applying both the seek and taxonomy modes.**

This file contains each type of information provided for each protein in the TXT and XLSX annotation files generated at the end of the run by *ProteoSeeker*. Each type of information belongs to a category and is specific to a field of information in that category. Each field of information in a category has a description. These fields and their information can be found in the TXT and XLSX annotation files. Each category may be repeated as many times as necessary for a protein depending on how many times this category of information has been collected for a protein (e.g., a protein may contain several domains, signal peptides and transmembrane regions). It should be noted that the categories, fields and descriptions of the information provided in the additional file are set in different order in the additional file compared to their order in the annotation files.

Additional file 4 TSV

**The Shannon index and the value of the filtering threshold computed automatically for non-gut and gut samples for each Kraken2 database and each sample.** Contains the Shannon index computed through KrakenTools for the predicted group of species by Kraken2, after abundance estimation from Bracken, for each database used by Kraken2 with the filtering thresholds computed for non-gut and gut samples, and for each sample during the taxonomy mode evaluation of *ProteoSeeker*. It also provides the filtering threshold computed by *ProteoSeeker*, for each Shannon index, for the case of non-gut and gut samples.

