## Supplemental_File_1 for "*ProteoSeeker*: A Feature-Rich Metagenomic Analysis Tool for Accessible and Comprehensive Metagenomic Exploration"

**ADDITIONAL FILE 1**

**Accessible and Comprehensive Metagenomic Exploration**

**1. Supplementary Text**

**1.1 Filtering a protein database**

The process of filtering a protein database according to certain protein names, is based on an independent Python command-line tool and module which was developed specifically for this process. This tool takes as input a set of file names and lists of protein names. Each file eventually will contain the proteins of the protein database which proteins contain in their headers at least one of the protein names of its corresponding list of protein names. This process has been parallelized. The protein database is divided into chunks. Each chunk is processed individually. The filtering takes place simultaneously for each chunk and is based on its list of protein names. This module is utilized by *ProteoSeeker* to filter the protein database when the seek mode and type 2 analysis or the taxonomy mode and the COMEBin/MetaBinner route is run, and the filtered database has not already been generated in a previous run.

**1.2 Metrics used in the evaluation of the taxonomy mode**

The metrics used in the evaluation of the Kraken2 and COMEBin/MetaBinner routes in the taxonomy mode are the following:

A. True Positive (TP) hits:

The number of common species between the predicted species by *ProteoSeeker* and the species which belong to the gold standard sample (“gold standard species”).

B. False Positive (FP) hits:

The number of species predicted by *ProteoSeeker* which species do not belong to the gold standard species.

C. False Negative (FN) hits:

The number of gold standard species that were not predicted by *ProteoSeeker*.

D. Sensitivity:

$$Sensitivity=\frac{TP}{TP + FN}$$

E. Precision:

$$Precision=\frac{TP}{TP + FP}$$

F. Accuracy:

$$Accuracy=\frac{TP + TN}{TP + TN + FP + FN}$$

G. F1 Score

$$F1 Score=2\frac{P * S}{P + S}=\frac{2 * TP}{2 * TP + FP + FN}$$

H. Jaccard Index

A: Set of gold standard species.

B: Set of predicted species by *ProteoSeeker*.

$$Jaccard Index=\frac{|A\cap B|}{|A\cup B|}=\frac{TP}{TP + FP + FN}$$

I. L1 norm

$$L1 norm=\sum_{i=1}^{n} |x_{i}-y_{i}|$$

where n is the number of species present in the set of gold standard species, $x_{i}$ is the relative abundance of the i^th^ species in the predicted profile, and $y_{i}$ is the relative abundance of the i^th^ species in the gold standard profile. The relative abundance of each predicted species, based on the COMEBin/MetaBinner route, is rounded up to two decimal places. The relative abundance of each predicted species, based on the Kraken2 route, is converted to a percentage and is rounded up to three decimal places based on the output of Bracken. A species that is not present in the predicted species takes a relative abundance equal to 0%.

**2. Supplementary Figures**

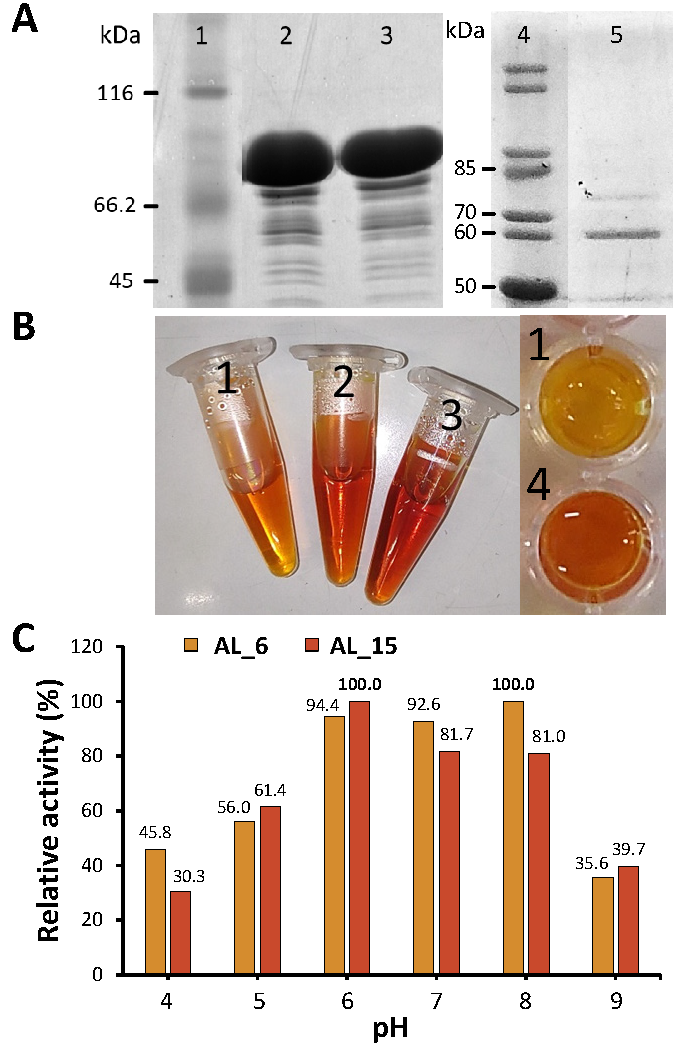

**Figure S1.** **Biochemical characterization of IMAC purified AL_6 and AL_15. (A)** SDS-PAGE analysis of AL_6 **(Lane 2)**, AL_15 **(Lane 3)** and AL_17 **(Lane 5)** amylases on a 15% polyacrylamide gel, with a protein standard in Lane 1 and Lane 4. **(B)** DNS assay (1) of the amylases at 70 °C for 1 hour (0.05% w/v final starch concentration). Samples include: No enzyme reaction **(1)**, AL_6 **(2)**, AL_15 **(3)**, and AL_17 **(4)**. During the amylolytic reaction, the reducing sugars produced react with 3,5-Dinitrosalicylic acid (DNS), forming 3-Aminosalicylic acid, which generates a distinct red color. The intensity of the red color indicates amylase activity. **(C)** pH-dependent activity of AL_6 and AL_15. Samples were incubated in buffers with pH ranging from 4 to 9, followed by DNS assay (0.05% w/v final starch concentration) to measure enzymatic activity via absorbance at 540 nm. The maximum absorbance values for AL_6 and AL_15 were set as 100% relative activity, and all other values were normalized to these maxima.

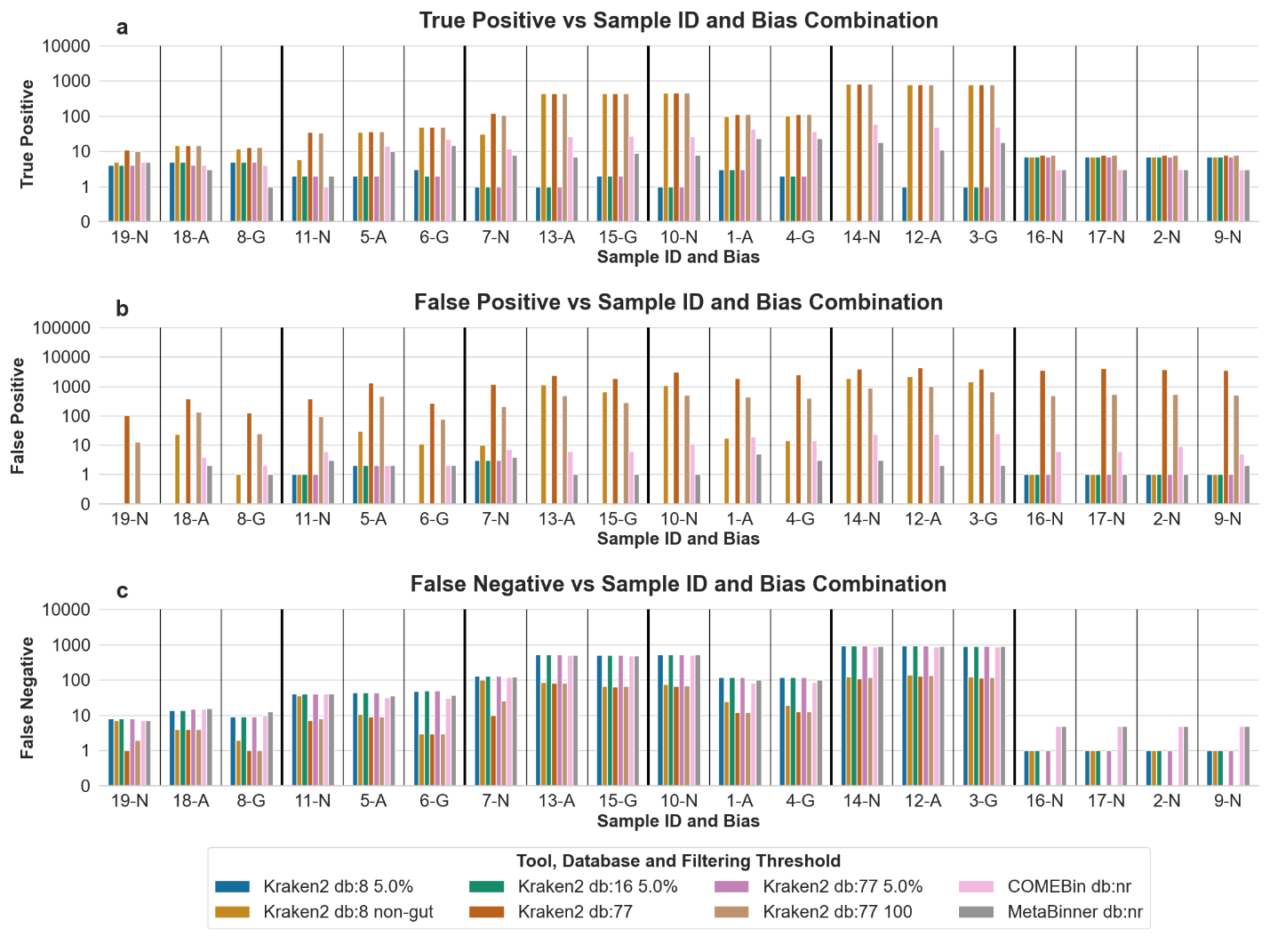

**Figure S2.** True positive **(a)**, false positive **(b)** and false negative **(c)** hits of the taxonomy mode evaluation results with the samples sorted based on their species-abundances and biases. The results were acquired by the ProteoSeeker runs for the selected combinations of Kraken2 databases and filtering thresholds plus the COMEBin/MetaBinner taxonomy methods for each sample of the 19 gold standard datasets. The selected combinations regarding the Kraken2 taxonomy route include the database of the Standard-8 collection with the filtering threshold of 5.0% (“Kraken2 db:8 5.0%”) and the non-gut filtering (“Kraken2 db:8 non-gut”), the database of the Standard-16 collection with the filtering threshold of 5.0% (“Kraken2 db:16 5.0%”) and the database of the Standard collection without a filtering threshold (“Kraken2 db:77”) and with the filtering thresholds of 5.0% (“Kraken2 db:77 5.0%”) and of 100 (“Kraken2 db:77 100”). The COMEBin/MetaBinner route was applied through COMEBin with the non-redundant protein database as the filtering target (“COMEBin db:nr”) and through MetaBinner with the non-redundant protein database as the filtering target (“MetaBinner db:nr”). The samples are sorted into groups of species-abundance. Samples 19, 18, 8 for 10 species from simulated reads, samples 11, 5, 6 for 40 species, samples 7, 13, 15 for 120 species, samples 10, 1, 4 for 500 species, samples 14, 12, 3 for 1000 species and samples 16, 17, 2, 9 for 10 species from cultures. The letters “N”, “A” and “G” on the labels stand for “No bias”, “AT-rich bias” and “GC-rich bias” respectively. The y-axis is in logarithmic scale (in base 10).

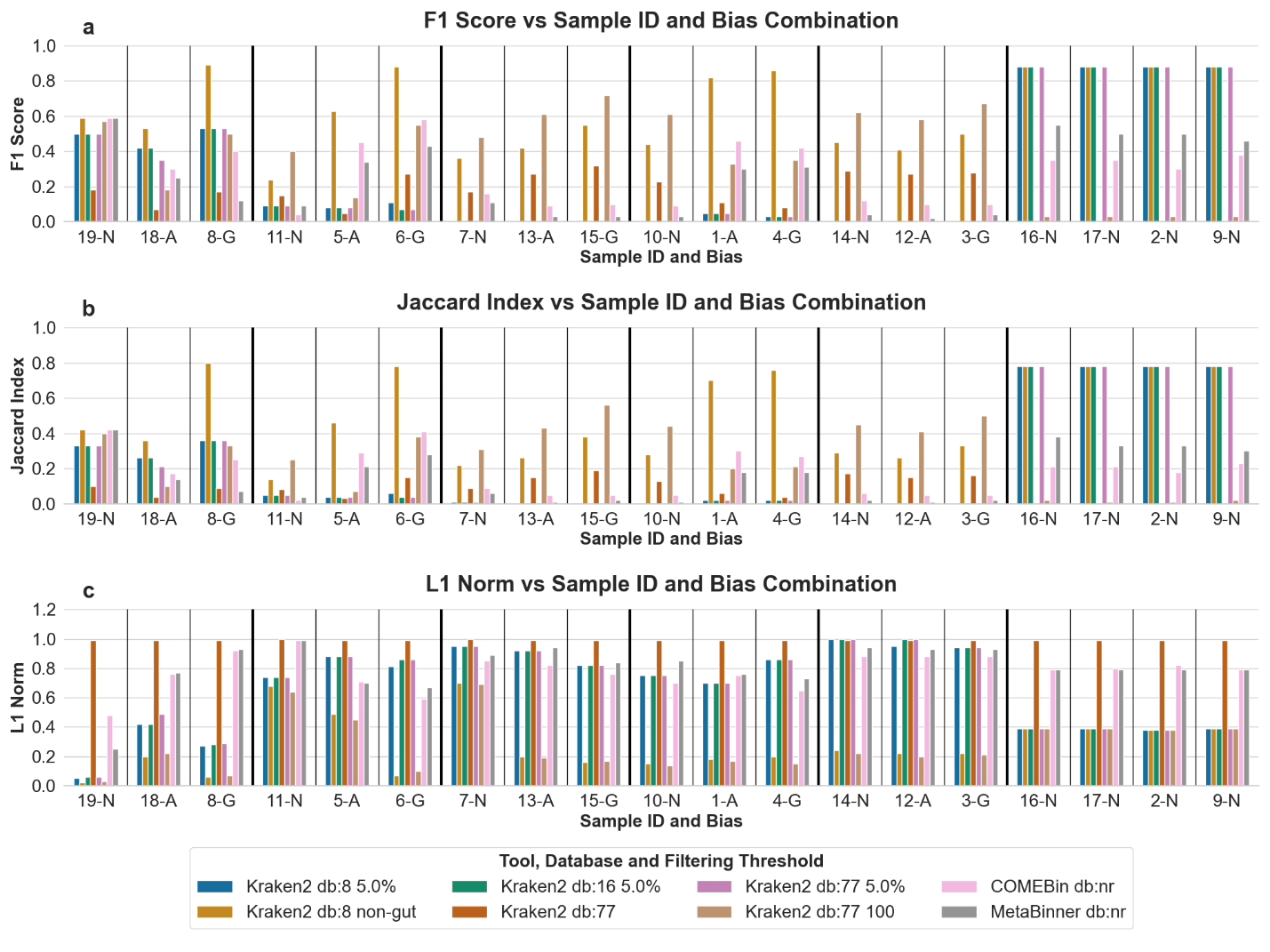

**Figure S3.** F1 score **(a)**, Jaccard index **(b)** and L1 norm **(c)** of the taxonomy mode evaluation results with the samples sorted based on their species-abundances and biases. The results were acquired by the ProteoSeeker runs for the selected combinations of Kraken2 databases and filtering thresholds plus the COMEBin/MetaBinner taxonomy methods for each sample of the 19 gold standard datasets. The selected combinations regarding the Kraken2 taxonomy route include the database of the Standard-8 collection with the filtering threshold of 5.0% (“Kraken2 db:8 5.0%”) and the non-gut filtering (“Kraken2 db:8 non-gut”), the database of the Standard-16 collection with the filtering threshold of 5.0% (“Kraken2 db:16 5.0%”) and the database of the Standard collection without a filtering threshold (“Kraken2 db:77”) and with the filtering thresholds of 5.0% (“Kraken2 db:77 5.0%”) and of 100 (“Kraken2 db:77 100”). The COMEBin/MetaBinner route was applied through COMEBin with the non-redundant protein database as the filtering target (“COMEBin db:nr”) and through MetaBinner with the non-redundant protein database as the filtering target (“MetaBinner db:nr”). The samples are sorted into groups of species-abundance. Samples 19, 18, 8 for 10 species from simulated reads, samples 11, 5, 6 for 40 species, samples 7, 13, 15 for 120 species, samples 10, 1, 4 for 500 species, samples 14, 12, 3 for 1000 species and samples 16, 17, 2, 9 for 10 species from cultures. The letters “N”, “A” and “G” on the labels stand for “No bias”, “AT-rich bias” and “GC-rich bias” respectively.

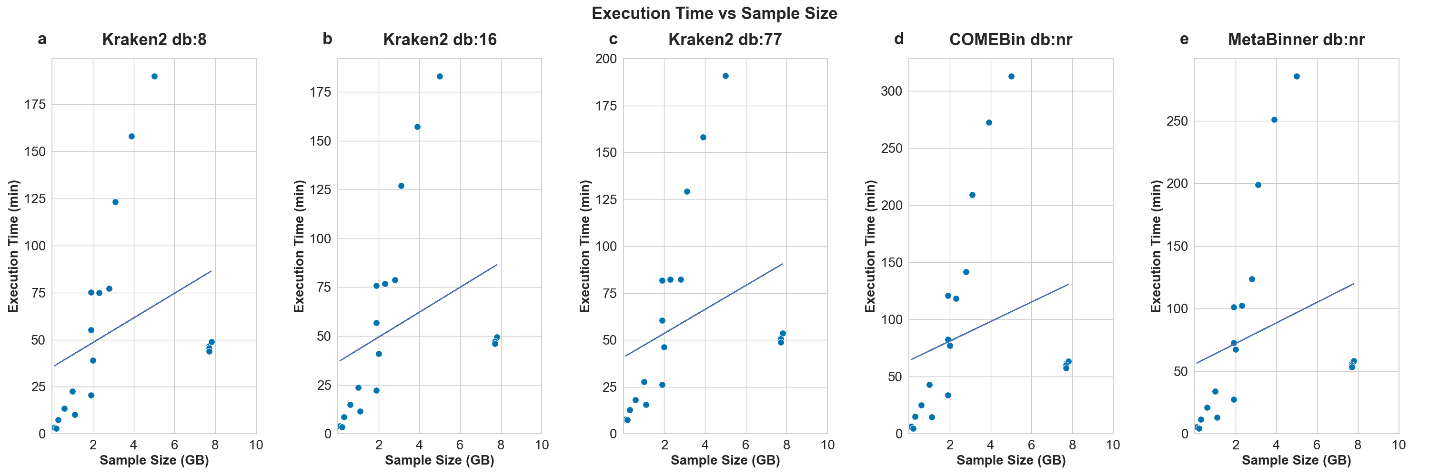

**Figure S4.** The total execution time of *ProteoSeeker*, based on its taxonomy mode evaluation, for each database and in turn for each of the 19 gold standard datasets, relative to their sizes. The databases used in the evaluation the Kraken2 route are **(a)** the Standard-8 collection, **(b)** the Standard-16 collection, **(c)** the Standard collection and for the COMEBin/MetaBinner route, **(d)** for COMEBin and for **(e)** MetaBinner the non-redundant (nr) database of NCBI. The size of a sample in GB, which equals the sum of the sizes of the paired-end FASTQ files extracted from the SRA dataset of the sample. In the case of the Kraken2 route, the effect of the filtering threshold is considered negligible as it may affect the number of species to be considered during the binning process and subsequent stages of the pipeline which effects in total account for very small changes in the execution time. The total execution time for each of the datasets does not include the time needed to download and process the SRA sample by *ProteoSeeker*. The initial stages of the pipeline, up to the stage of gene prediction, have the exact same execution time for the different runs of the same sample. A straight line has been fitted to the data points of each group of runs based on the combination of the taxonomy route and database.

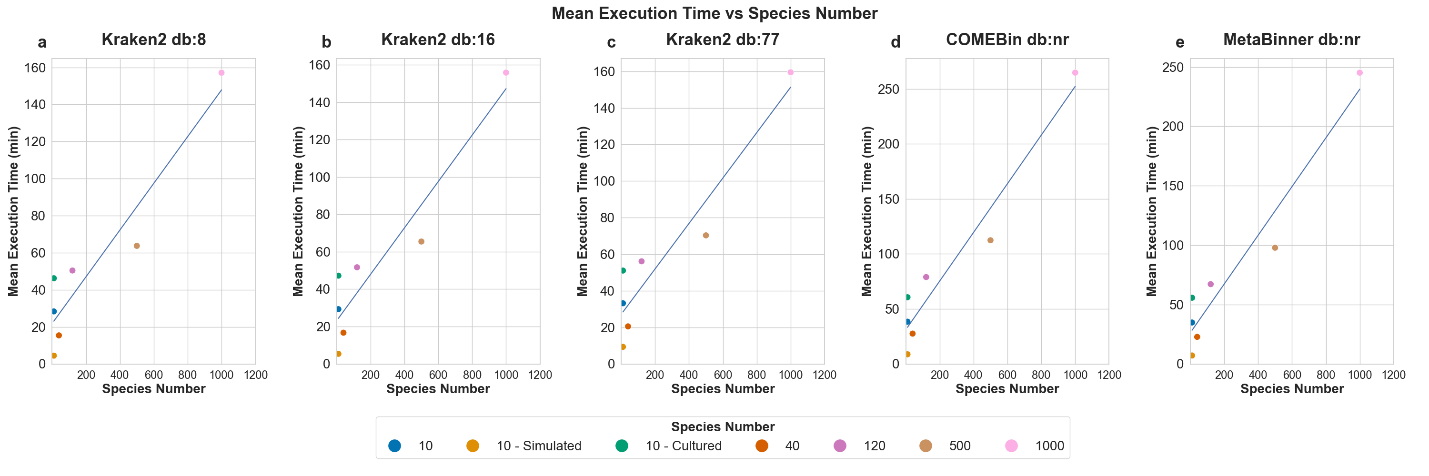

**Figure S5.** The mean execution time of *ProteoSeeker*, based on its taxonomy mode evaluation, for each database and in turn for each of the 19 gold standard datasets, relative to their species abundance. The databases used in the evaluation the Kraken2 route are **(a)** the Standard-8 collection, **(b)** the Standard-16 collection, **(c)** the Standard collection and for the COMEBin/MetaBinner route, **(d)** for COMEBin and for **(e)** MetaBinner the non-redundant (nr) database of NCBI. In the case of the Kraken2 route, the effect of the filtering threshold is considered negligible as it may affect the number of species to be considered during the binning process and subsequent stages of the pipeline which effects in total account for very small changes in the execution time. For the group of 10 species the mean execution time has also been computed for the two subgroups of 10 species based on whether they originate from simulated reads or from cultures. Each of the total execution times based on which the mean times were computed does not include the time needed to download and process the SRA sample by *ProteoSeeker*. The initial stages of the pipeline, up to the stage of gene prediction, have the exact same execution time for the different runs of the same sample. A straight line has been fitted to the data points of each group of runs based on the combination of the taxonomy route and database.

**3. Supplementary Tables**

**Table S1.** Information about each of the 19 gold standard datasets. For each dataset its sample ID, its SRA code, its species number, its category, its bias and the size of the paired-end FASTQ files extracted from the SRA file and analyzed by *ProteoSeeker* are provided.

| **Sample ID** | **SRA code** | **Species number** | **Category** | **Bias** | **Size (GB)** |
| --- | --- | --- | --- | --- | --- |
| 8 | SRR12829159 | 19 | 10 | GC-rich bias | 0.1297 |
| 19 | SRR12829170 | 12 | 10 | No bias | 0.2049 |
| 18 | SRR12829169 | 14 | 10 | AT-rich bias | 0.3474 |
| 17 | SRR12829162 | NA | 10 | No bias | 7.7 |
| 16 | SRR12829163 | NA | 10 | No bias | 7.7 |
| 9 | SRR12829160 | NA | 10 | No bias | 7.8 |
| 2 | SRR12829161 | NA | 10 | No bias | 7.7 |
| 6 | SRR12829156 | 46 | 40 | GC-rich bias | 0.6313 |
| 5 | SRR12829157 | 52 | 40 | AT-rich bias | 0.9808 |
| 11 | SRR12829158 | 42 | 40 | No bias | 1.1 |
| 7 | SRR12829155 | 132 | 120 | No bias | 1.9 |
| 13 | SRR12829154 | 124 | 120 | AT-rich bias | 1.9 |
| 15 | SRR12829153 | 124 | 120 | GC-rich bias | 1.9 |
| 1 | SRR12829168 | 508 | 500 | AT-rich bias | 2.0 |
| 10 | SRR12829152 | 536 | 500 | No bias | 2.3 |
| 4 | SRR12829167 | 528 | 500 | GC-rich bias | 2.8 |
| 3 | SRR12829164 | 934 | 1000 | GC-rich bias | 3.1 |
| 14 | SRR12829166 | 949 | 1000 | No bias | 3.9 |
| 12 | SRR12829165 | 912 | 1000 | AT-rich bias | 5.0 |

**Table S2.** Information about the different evaluation cases of the taxonomy mode (or taxonomy routes) of *ProteoSeeker*. For each case, the tool utilized by *ProteoSeeker* in the taxonomy route, and the database utilized in the evaluation are provided. Each gold standard sample was run through each of these 5 combinations of taxonomy route, tool and database.

| **Taxonomy route** | **Tool** | **Database** |
| --- | --- | --- |
| Kraken2 | Kraken2 | Kraken 2 / Bracken Refseq index, Collection: Standard-8 |
| Kraken2 | Kraken2 | Kraken 2 / Bracken Refseq index, Collection: Standard-16 |
| Kraken2 | Kraken2 | Kraken 2 / Bracken Refseq index, Collection: Standard |
| COMEBin/MetaBinner | COMEBin | NCBI, non-redundant (nr) database |
| COMEBin/MetaBinner | MetaBinner | NCBI, non-redundant (nr) database |

**Table S3.** Information for the taxonomy classification of experimentally verified and studied enzymes CA-KR1, CA_201 and AL_17, discovered by the seek mode of *ProteoSeeker*. For each enzyme the species of the hit with the lowest E-value acquired from running BLASTP against the nr database is provided. For the Kraken2 and COMEBin/MetaBinner taxonomy route, the taxonomy classification of the enzyme is provided. If the taxonomy classification could not be inferred for the protein, then “None” is noted. For the Kraken2 taxonomy route three databases were used for the analysis, the Kraken 2 / Bracken Refseq indexes of the Standard-8 (“Kraken2 db:8”), Standard-16 (“Kraken2 db:16”) and Standard (“Kraken2 db:77”) collections. The COMEBin/MetaBinner route was applied based on the non-redundant (nr) database of NCBI through COMEBin (“COMEBin db:nr”) and through MetaBinner (“MetaBinner db:nr”). No filtering threshold was applied to the abundances or relative abundances of the taxa predicted in each taxonomy route.

| **Tool and Database** | **CA-KR1** | **CA_201** | **AL_17** |
| --- | --- | --- | --- |
| COMEBin db:nr | *Pyrobaculum aerophilum* | *Novosphingobium*  *Novosphingobium* sp. *AAP1*  *Novosphingobium* sp. *BK256*  *Novosphingobium* sp. *BK280*  *Novosphingobium* sp. *BK258*  *Novosphingobium* sp. *BK267*  *Novosphingobium* *pokkalii* | *Rheinheimera* sp. *MM224* |
| MetaBinner db:nr | None | *Novosphingobium*  *Novosphingobium* sp. *AAP1*  *Novosphingobium* sp. *BK256*  *Novosphingobium* sp. *BK280*  *Novosphingobium* sp. *BK258*  *Novosphingobium* sp. *BK267*  *Novosphingobium* *pokkalii* | *Rheinheimera* sp. *MM224* |
| Kraken2 db:8 | None | *Novosphingobium humi* | *Rheinheimera mangrovi* |
| Kraken2 db:16 | None | *Novosphingobium humi* | *Rheinheimera mangrovi* |
| Kraken2 db:77 | *Moraxella bovoculi* | *Novosphingobium humi* | *Rheinheimera* sp. *MM224* |
| BLASTP db:nr | *Pyrobaculum aerophilum* | *Novosphingobium* sp. | *Rheinheimera mesophila* |
